# Detection of Cytosolic Ion Concentrations by the Trans-Golgi Network/Early Endosome is Important for Salt Tolerance

**DOI:** 10.1101/2025.08.13.670069

**Authors:** Daniel McKay, Upendo Lupanga, Michelle Uebele Perez, Melanie Krebs, Stefanie Wege, Michael Grabe, Karin Schumacher

**Affiliations:** Cell Biology, Centre for Organismal Studies (COS), Heidelberg University, Im Neuenheimer Feld 230, 69120 Heidelberg, Germany; INRES, Plant Nutrition, University of Bonn, Germany; Cardiovascular Research Institute and Department of Pharmaceutical Chemistry, University of California, San Francisco, United States

## Abstract

Plant survival requires cellular sensing systems that detect and respond to nutrient and ion fluctuations. The ability to monitor and adjust solute concentrations is essential for managing environmental stresses, particularly salt stress, yet the molecular mechanisms underlying cellular ion sensing remain largely unknown. Here, we combine luminal pH measurements with mathematical modelling of ion transport to investigate the role of the Trans-Golgi Network/Early Endosome (TGN/EE) for cellular ion homeostasis. Our results demonstrate that cytosolic concentrations of Cl^-^, Na^+^ and K^+^ directly impact ion transport activity at the TGN/EE, thereby influencing luminal pH dynamics. Specifically, we find that the TGN/EE lumen becomes more alkaline when plants are exposed to elevated NaCl concentrations, indicating a previously unrecognised role for endomembrane pH regulation in salt stress responses. Furthermore, we demonstrate that TGN/EE ion transport mutants that fail to exhibit NaCl-induced alkalisation display hypersensitivity to Na^+^. These findings indicate that the translation of elevated cytosolic ion concentrations into TGN/EE luminal alkalisation, represents an important mechanism for conferring Na^+^ tolerance in plant cells.

## Introduction

Ion transport in plants is vital for nutrition, cell growth, signalling and response to environmental changes and stresses such as salt stress. Coordination of ion transport across multiple cells types allows the partitioning of ions in plants to provide valuable nutrients where they are required and prevent the accumulation of detrimental ions like Na^+^ in sensitive tissues. Similar to organism-wide coordination of ion fluxes, individual cells must balance transport across numerous internal membranes and compartments. Recent advances have enabled the characterisation of ion concentrations in larger cellular compartments such as the vacuole, nucleus and cytosol (Ramakrishna et al., 2025). However, direct measurement of ion concentrations within endomembrane compartments remains challenging due to their small sizes and biochemical environments. Conventional mass spectrometry analytical techniques lack spatial resolution (Zhang et al., 2024), while application of fluorescent indicators can be limited due to difficulties in subcellular targeting, inappropriate dissociation constants, or reduced brightness due to the acidic environments of endomembrane compartments (Sadoine et al., 2021; Shinoda et al., 2018; Suzuki et al., 2016). These technical constraints have resulted in a knowledge gap in understanding the mechanisms of cellular ion homeostasis across different cellular compartments. In the absence of reliable direct measurements, mathematical modelling offers a powerful approach to infer ion dynamics and to bridge uncertainties arising from limited experimental data. By integrating physiological, biochemical, and imaging information, these models not only generate testable predictions but also yield deeper insights into ion regulation within endomembrane compartments.

One such critical endomembrane compartment, is the trans-Golgi Network/Early Endosome (TGN/EE), which coordinates vesicular trafficking in plant cells. Several ion transport proteins within the TGN/EE are crucial for its function in directing vesicular cargo to specific cellular destinations (Dragwidge et al., 2019; Luo et al., 2015; McKay et al., 2022). Disruption of TGN/EE-mediated trafficking results in multiple phenotypic defects reflecting the various cellular processes that depend on this essential trafficking pathway (Rosquete et al., 2018). The ion transporters localised to the TGN/EE regulate its ion composition, which can be quantified as changes to the luminal pH. The TGN/EE lumen typically maintains a pH of approximately 5.6, acidified by the activity of the vacuolar H^+^-ATPase (V-ATPase) complex (Luo et al., 2015). The V-ATPase complex exhibits dual localisation, functioning at both the TGN/EE and the vacuolar membrane (tonoplast), with its TGN/EE localisation mediated by the incorporation of the subunit isoform VHA-a1 (Dettmer et al., 2006). The proton gradient established by V-ATPase drives influx of Cl^-^ through Cl^-^/H^+^ antiporters CLCd and CLCf, as well as influx of K^+^ through the K^+^/H^+^ antiporters NHX5 and NHX6 (Bassil et al., 2011; Dragwidge et al., 2019, Dragwidge et al., 2018; Fecht-Bartenbach et al., 2007; Reguera et al., 2015; Scholl et al., 2021). The accumulated K^+^ and Cl^-^ ions are subsequently exported from the TGN/EE through the Cl^-^/K^+^ symporter, CCC1, thereby completing the ion transport circuit (Figure 1A; Colmenero-Flores et al., 2007; Han et al., 2020; Henderson et al., 2015; McKay et al., 2022). Mutations that disrupt pH homeostasis within the TGN/EE, such as those observed for the V-ATPase mutant *det3* (Schumacher et al., 1999), the *nhx5 nhx6* double mutant (Reguera et al., 2015) and the *ccc1* mutant (Colmenero-Flores et al., 2007) lead to a general reduction in bulk vesicular trafficking resulting in a range of growth defects (Bassil et al., 2011; Dettmer et al., 2006; Dragwidge et al., 2019, Dragwidge et al., 2018; Han et al., 2020; McKay et al., 2022). Remarkably, the resulting phenotypes differ among these mutants, suggesting that specific alterations in TGN/EE ion homeostasis result in distinct changes in trafficking patterns. Single knockout mutants of *clcd, clcf, nhx5*, or *nhx6* show no changes to luminal pH or vesicular trafficking, thus maintaining wild-type-like growth. Notably, while the *nhx5 nhx6* double knockout is viable, though showing severe growth defects, the *clcd clcf* double knockout is embryo lethal (Bassil et al., 2011; Reguera et al., 2015; Scholl et al., 2021), which suggests at least partially redundant functions within these transporter families. While the two TGN/EE-resident NHX transporters in Arabidopsis likely originated from a gene duplication event (Pires et al., 2013), CLCd and CLCf belong to different CLC subclades, with CLCf likely being of plastidial origin, indicating that CLCd and CLCf have independently acquired roles in TGN/EE luminal ion regulation (Scholl et al., 2021). Although these transporters share the same subcellular localization, distinct and non-redundant functions, such as a role in salt stress responses, have not yet been reported.

**Figure 1.**
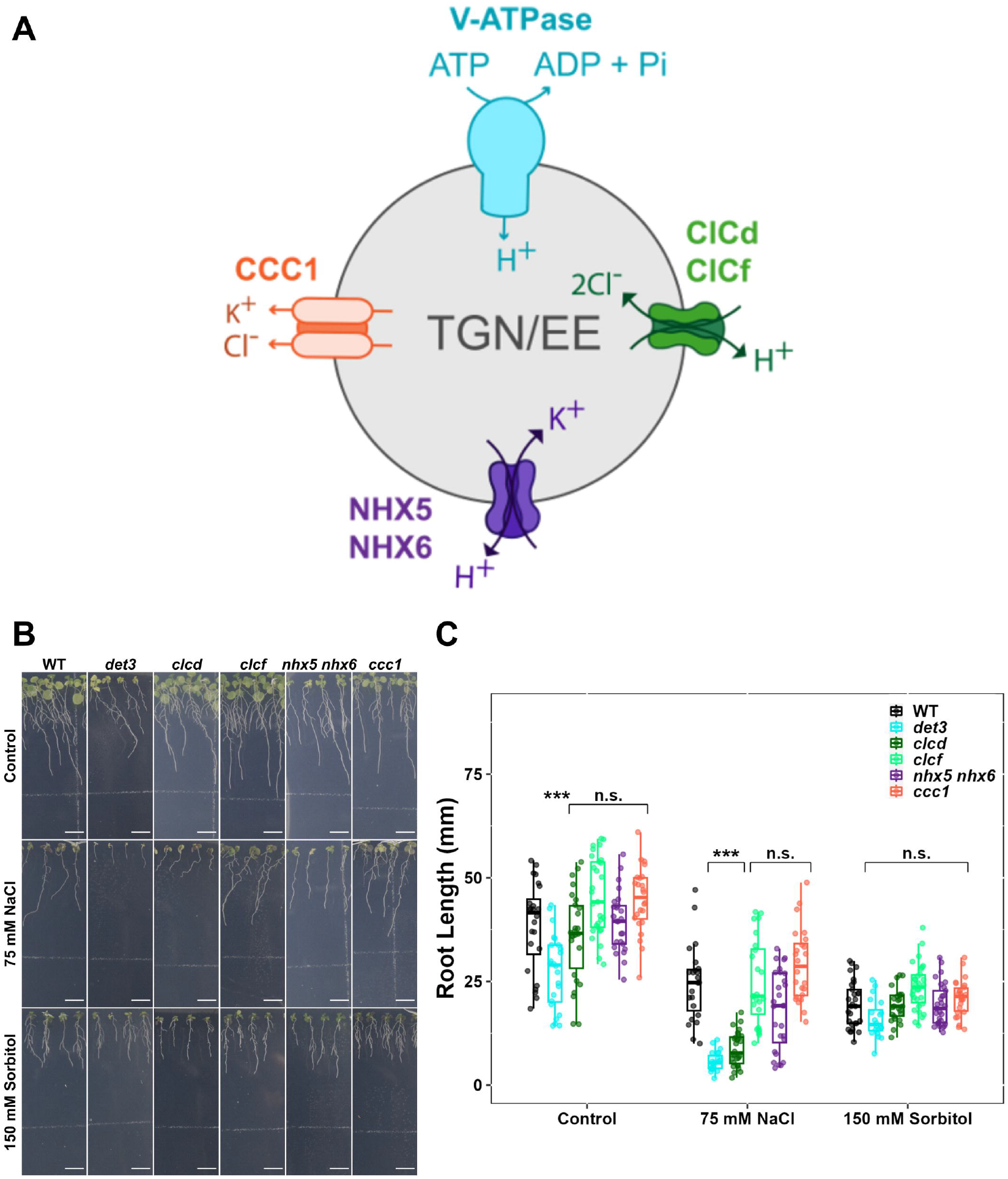
Specific TGN/EE ion transporters have a role in tolerance to salt stress. **A)** A representation of the TGN/EE ion transport circuit including the proton pumping V-ATPase, the 2Cl^-^/H^+^ antiporters CLCd and CLCf, the K^+^/H^+^ antiporters NHX5 and NHX6 and the Cl^-^/K^+^ symporter CCC1. **B-C)** The primary root length of *det3, nhx5/nhx6* and *ccc1* mutants grown constitutively on 75 mM NaCl is shorter than WT plants. No genotypes show defects in primary root growth on the osmotic stress, 150 mM sorbitol, comparative to WT. The upper white line in the background of images is part of the equipment. Scale bars = 10 mm. Statistical significance was determined via one-way ANOVA, comparing mutant genotypes to WT under the same conditions. P values are indicated by***<0.001<**<0.01<*<0.05<n.s. For the boxplots, centre line = median, box limits = upper and lower quartiles, whiskers = range to a maximum of 1.5× the interquartile range, points = individual data points.

Hypersensitivity to elevated NaCl levels has been observed in TGN/EE ion transport mutants with impaired V-ATPase activity. For instance, the *det3* mutant exhibits severely reduced root growth in response to 100 mM NaCl, while inducible RNAi knockdown lines targeting the V-ATPase subunit VHA-a1 show impaired root development even at 25 mM NaCl (Batelli et al., 2007; Krebs et al., 2010). Similarly, simultaneous knockout of the TGN/EE-resident K^+^/H^+^ antiporters *NHX5* and *NHX6* leads to defects in growth and germination on media containing 100 mM NaCl (Bassil et al., 2011). Given the multifaceted phenotypes of these mutants and the severity of their growth defects, such salt hypersensitivity might reasonably be attributed to indirect effects from disruptions in other cellular processes. Although the *ccc1* mutant also displays disrupted TGN/EE pH, defective trafficking, and marked growth defects, but notably does not exhibit hypersensitivity to elevated salt, thereby challenging the notion that impaired TGN/EE function alone is sufficient to explain salt sensitivity in these mutants (Henderson et al., 2015). Together, these observations suggest that salt hypersensitivity in TGN/EE ion transport mutants cannot be ascribed solely to general disruptions in TGN/EE function, but rather suggest a more specific interaction between TGN/EE ion transport activity and salt stress response.

Ion transporters are essential for salinity tolerance because they maintain ionic balance, and cellular osmolality, thereby protecting cells from Na^+^ toxicity (Choudhary et al., 2023; Zelm et al., 2020). Excess amounts of Na^+^ disrupts cytosolic K^+^ homeostasis, which is critical to maintain normal cellular functions (Benito et al., 2014). Plants counter elevated Na^+^ levels through the coordinated action of multiple classes of transport proteins, which actively export Na^+^ from the cytosol to the apoplast, sequester it into vacuoles, retrieve it from the xylem of root tissues to prevent translocation to sensitive aerial parts and selectively load it into the phloem of shoot tissues to facilitate its redistribution to the roots (Wegner et al., 2025; Yang and Guo, 2018; Zelm et al., 2020; Zhou et al., 2024). These adaptive responses to salt stress are thought to act downstream of cellular signalling events, that include an immediate increase in cytosolic Ca^2+^, alterations in cellular pH and increased production of reactive oxygen species (ROS; Choi et al., 2014; Jiang et al., 2019; Steinhorst et al., 2022; Yang and Guo, 2018; Zelm et al., 2020). Beyond their acknowledged roles as downstream targets modulated by NaCl-induced sensing and signaling, ion transport networks have themselves been proposed as primary sensors of cellular ion and nutrient status (Contador-Álvarez et al., 2025; Dreyer et al., 2022; Wegner et al., 2025). Such a model predicts, that transport networks can directly translate fluctuations in extracellular ion and nutrient concentrations into intracellular pH shifts and Ca^2+^ signals thereby initiating the signalling cascade that trigger adaptive downstream responses (Dreyer et al., 2022).

Here, we combine luminal pH measurements with mathematical modelling to investigate the role of the TGN/EE in cellular ion homeostasis and its interaction with salt stress. We demonstrate that TGN/EE ion transport, specifically the activity of V-ATPase, NHXs, and CLCd, but not CLCf and CCC1, is crucial for tolerance to Na^+^ stress, thereby also identifying a previously unrecognised function of CLCd. We find that NaCl induces alkalinisation of the TGN/EE, and this alkalinisation depends on the ion transport network of the TGN/EE. Plants unable to alkalinise their TGN/EE in response to NaCl exhibit Na^+^ hypersensitivity. Using a mathematical model of the TGN/EE ion transport network, we show that TGN/EE alkalinisation is a direct result of elevated cytosolic NaCl concentrations, which alter the dynamics of the TGN/EE ion transport circuit. Importantly, this shift in pH does not require additional receptors, kinases or Na^+^ binding domains to mediate the response. We further reveal that TGN/EE alkalinisation occurs in a dose-dependent manner, with pH increasing proportionally to cytosolic NaCl concentrations. Notably, this response is not specific to Na^+^ and can also be triggered by K^+^ and Cl^-^. Therefore, we propose that the TGN/EE ion transport circuit serves as a detector for cytosolic ion concentrations, transducing ionic changes into luminal pH adjustments that are essential for cellular tolerance to high Na^+^ concentrations. Our work thus advances the understanding of plant salt stress responses by identifying a novel mechanism of cellular stress detection.

## Results

### TGN/EE localised ion transport proteins are important for tolerance to Na^+^ stress

Multiple studies have demonstrated that mutants defective in TGN/EE ion transport proteins exhibit salt hypersensitivity, suggesting a link between TGN/EE ion homeostasis and salt tolerance mechanisms (Bassil et al., 2011; Batelli et al., 2007; Krebs et al., 2010). To investigate the specific role of TGN/EE ion homeostasis in plant salt sensitivity, we systematically determined which TGN/EE transport proteins contribute to salt stress tolerance. The primary root length of wild-type (WT), *det3, clcd, clcf, nhx5 nhx6*, and *ccc1* plants was measured after 14 days of growing either on control media or medium containing 75 mM NaCl. Compared to WT plants, *det3, clcd*, and *nhx5 nhx6* plants displayed a more severe reduction of root length under salt stress conditions, newly identifying *clcd* knockouts as salt hypersensitive and confirming the previously reported salt hypersensitivity of *det3* and *nhx5 nhx6* mutants (Figure 1B-C). Among the tested mutants, *clcd* displayed the most pronounced hypersensitivity. In contrast, the root length of *clcf* and *ccc1* mutants was similar to WT under NaCl stress, consistent with previous findings showing no salt hypersensitivity in *clcf* mutants (McKay et al., 2025). Under non-stress growth conditions, CLCd and CLCf function redundantly in TGN/EE pH regulation, as both single mutants exhibit wild-type phenotypes whereas the double mutant exhibits embryo lethality (Scholl et al., 2021). However, here we reveal that CLCd serves a critical non-redundant role during salt stress that CLCf cannot compensate for. In contrast to other TGN/EE ion transport mutants, such as *det3* and *nhx5 nhx6* which display general growth defects (Bassil et al., 2011; Schumacher et al., 1999), *clcd* mutants maintain growth indistinguishable from WT plants under control conditions (Figure S1; Scholl et al., 2021). This distinction suggests that the salt hypersensitivity observed in TGN/EE ion transport mutants is not merely a secondary consequence of their general growth defects but rather points to a specific role for TGN/EE ion homeostasis during salt stress.

To determine whether the salt hypersensitivity observed in *det3, nhx5 nhx6* and *clcd* is due to osmotic or ionic effects of salt treatment, we measured the primary root length of plants grown on medium containing 150 mM sorbitol, which imposes osmotic stress equivalent to that of 75 mM NaCl. Under these conditions, all transport mutants displayed root lengths comparable to WT, indicating that the observed hypersensitivity to salt is not due to compromised tolerance to osmotic stress (Figure 1B-C). To identify which ionic component of NaCl contributes to hypersensitivity in TGN/EE transport mutants under salt stress, primary root length of seedlings grown on media without Na^+^ but containing a mixture of 75 mM Cl^-^ salts (25 mM KCl, 12.5 mM MgCl_2_ and 12.5 mM CaCl_2_) was measured. With this treatment, *det3* mutants still exhibited hypersensitivity, whereas *clcd* and *nhx5 nhx6* knockout plants were not hypersensitive. This suggests that Na^+^, but not Cl^-^, is a causative factor responsible for hypersensitivity in *clcd* and *nhx5 nhx6* mutants during NaCl stress (Figure 2A-B). We next tested whether Cl^-^ contributes to hypersensitivity in TGN/EE transport mutants. We measured primary root length when seedlings were grown on media containing 75 mM Na^+^ without additional Cl^-^, using either 75 mM NaNO_3_ or 37.5 mM Na_2_SO_4_. Under both treatments, *det3, clcd*, and *nhx5 nhx6* exhibited hypersensitivity towards salt stress (Figure 2C-D). This confirms, that Cl^-^ is not required for salt hypersensitivity in any TGN/EE transport mutant tested. In summary, *nhx5 nhx6* and *clcd* plants are specifically hypersensitive to Na^+^, while *det3* exhibits hypersensitivity to either both Na^+^ and Cl^-^, or is hypersensitive to one of the other cations present in the Cl^-^ salts treatment.

**Figure 2.**
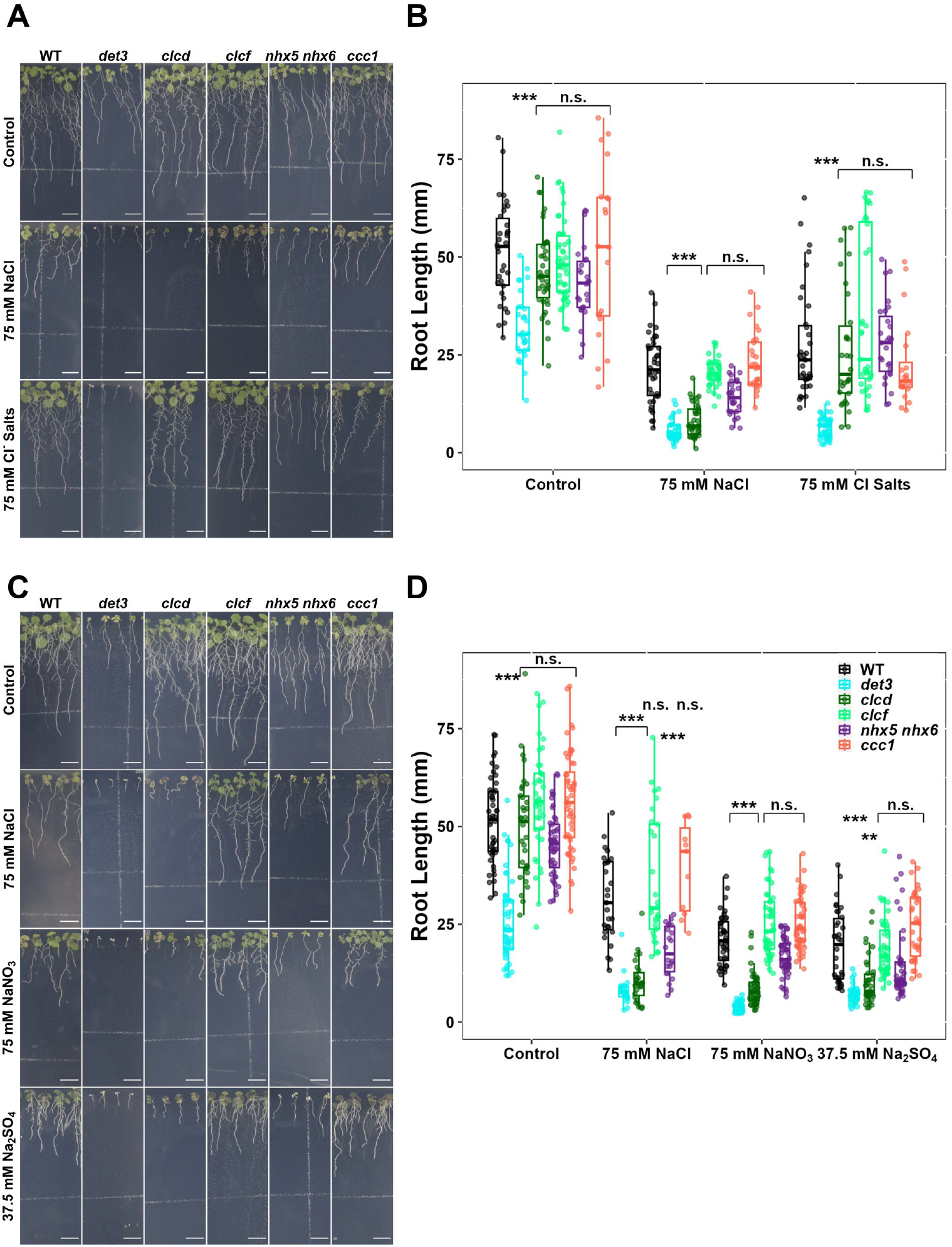
The TGN/EE ion transporter circuit has a role specifically under Na^+^ stress. **A-B)** *det3* but not *nhx5/nhx6* or *clcd* have defects in primary root length in response to a Cl-salt mixture containing 25 mM KCl, 12.5 mM MgCl2 and 12.5 mM CaCl2. **C-D)** Primary root length of *det3, nhx5/nhx6* and *clcd* plants is shorter comparative to WT when grown on media containing 75 mM NaNO_3_ or 37.5 mM Na_2_SO_4_. For all experiments, plants were imaged at 14 days old. The upper white line in the background of images is part of the equipment. Scale bars = 10 mm. Statistical significance was determined via one-way ANOVA, comparing mutant genotypes to WT under the same conditions. P values are indicated by ***<0.001<**<0.01<*<0.05<n.s. For the boxplots, centre line = median, box limits = upper and lower quartiles, whiskers = range to a maximum of 1.5× the interquartile range, points = individual data points.

To confirm the importance of TGN/EE ion transporters during Na^+^ stress beyond the seedling stage and to specifically validate the newly identified role of CLCd, we conducted growth experiments using WT, *clcd*, and *clcf* mutant plants. Plants were grown for four weeks in hydroponic culture with or without 75 mM NaNO_3_. Biomass analyses revealed that *clcd* mutants exhibited more severe growth defects in response to NaNO_3_ treatment than WT or *clcf* plants. This validates the critical role CLCd plays in mediating tolerance to Na^+^ stress across multiple tissues and developmental stages (Figure S1).

Additionally, we examined the subcellular localisation of fluorescent protein fusions of VHAa1, CLCd, CLCf, NHX6, and CCC1 during salt stress. Localisation analyses was performed in root epidermal cells of seedlings grown either under control conditions or in the presence of 75 mM NaCl, a concentration sufficient to induce salt stress-mediated growth inhibitions. Under control conditions, all proteins exhibit the typical punctate TGN/EE localisation pattern previously described (Figure 3; Bassil et al., 2011; Dettmer et al., 2006; Drakakaki et al., 2012; Fecht-Bartenbach et al., 2007; Groen et al., 2014; Henderson et al., 2015; McKay et al., 2025, 2022; Scholl et al., 2021). This punctate pattern was unchanged in roots grown in the presence of 75 mM NaCl stress. Therefore, the salt hypersensitivity observed in *clcd* and *nhx5 nhx6* mutants is due to loss of their transport activity in the TGN/EE.

**Figure 3.**
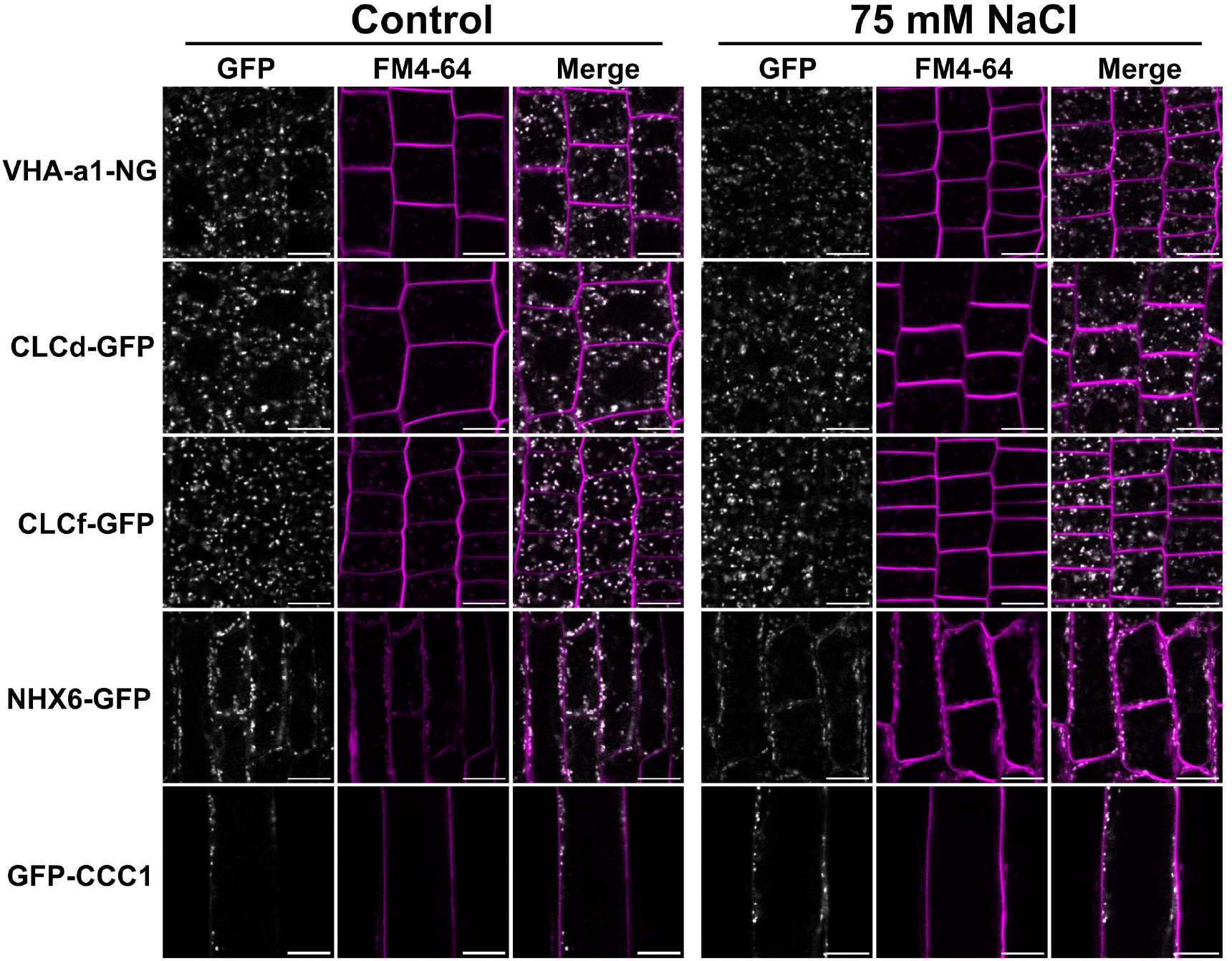
TGN/EE ion transport is important for tolerating Na^+^ stress. CLSM analysis shows that VHA-a1 (of the V-ATPase complex), CLCd, CLCf, NHX6 and CCC1 tagged with GFP or neongreen localise to small TGN/EE like puncta, where all proteins have previously been localised to, when grown on standard 1/2 MS media or media containing an additional 75 mM NaCl. Root epidermal cells of the elongation zone for VHAa1, CLCd, CLCf and NHX6 or trichoblast cells in mature roots were imaged in plants were imaged at 9 days old. Counterstaining to mark the PM was performed using 25 µM FM4-64 for 10 minutes. Images are single layers taken from the cortical zone of the cell. Scale bars = 10 um.

### CLCd function is not limiting for bulk cargo trafficking under salt stress conditions

To determine whether compromised TGN/EE-mediated trafficking underlies the Na^+^ hypersensitivity in TGN/EE ion transport mutants, we investigated whether CLCd loss-of-function causes cellular cargo trafficking defects during exposure to NaCl stress. The *clcd* mutant was selected for analysis as it exhibits normal vesicular trafficking under non-stress conditions (Scholl et al., 2021), enabling discrimination between general and Na^+^-induced trafficking defects. The *ccc1* mutant served as positive control, given its established defects in secretory and endocytic trafficking (McKay et al., 2022). Secretory and endocytic trafficking was analysed using established markers secRFP (Zheng et al., 2005) and spRFP (Hunter et al., 2007), which report bulk cargo trafficking from the endoplasmic reticulum (ER) to the apoplast and from the ER to vacuole, respectively. To measure efficacy of cargo trafficking, fluorescence microscopy analysis was performed in epidermal root cells and trafficking efficacy was expressed as a ratio between apoplast-to-ER-signal or vacuole-to-ER signal, respectively. Under control conditions, both markers showed varying degrees of intracellular ER signal. Under control conditions, secretory trafficking in *clcd* was unaffected, consistent with previous findings that CLCd is dispensable for trafficking under non-stressed conditions (Scholl et al., 2021). Salt stress did not result in inhibition of secretory trafficking in WT or *clcd* plants. Conversely, in *ccc1* mutants which were previously reported to have impaired secretory trafficking, salt stress resulted in decreased secretory trafficking (Figure 4A-B; McKay et al., 2022). Together, these results demonstrate that a reduction of bulk secretory trafficking is not the cause of Na^+^ hypersensitivity in TGN/EE transporter mutants as Na^+^ does not perturb secretory trafficking in the Na^+^ hypersensitive mutant *clcd* but does impact trafficking in *ccc1*, which is not hypersensitive to Na^+^. Under control conditions, bulk cargo trafficking to the vacuole was greater in *clcd* compared to WT while being slightly reduced in *ccc1* compared to WT, as previously reported (McKay et al., 2022). Salt stress markedly decreased vacuolar trafficking in WT plants, but this decrease was less pronounced in *clcd* mutants and absent in *ccc1* mutants (Figure 4C-D). Taken together, these results demonstrate that secretory and endocytic trafficking of bulk cargo in *clcd* is not affected by Na^+^ to a greater degree than in WT, suggesting that CLCd is not critical for maintaining bulk protein trafficking during salt stress conditions.

**Figure 4.**
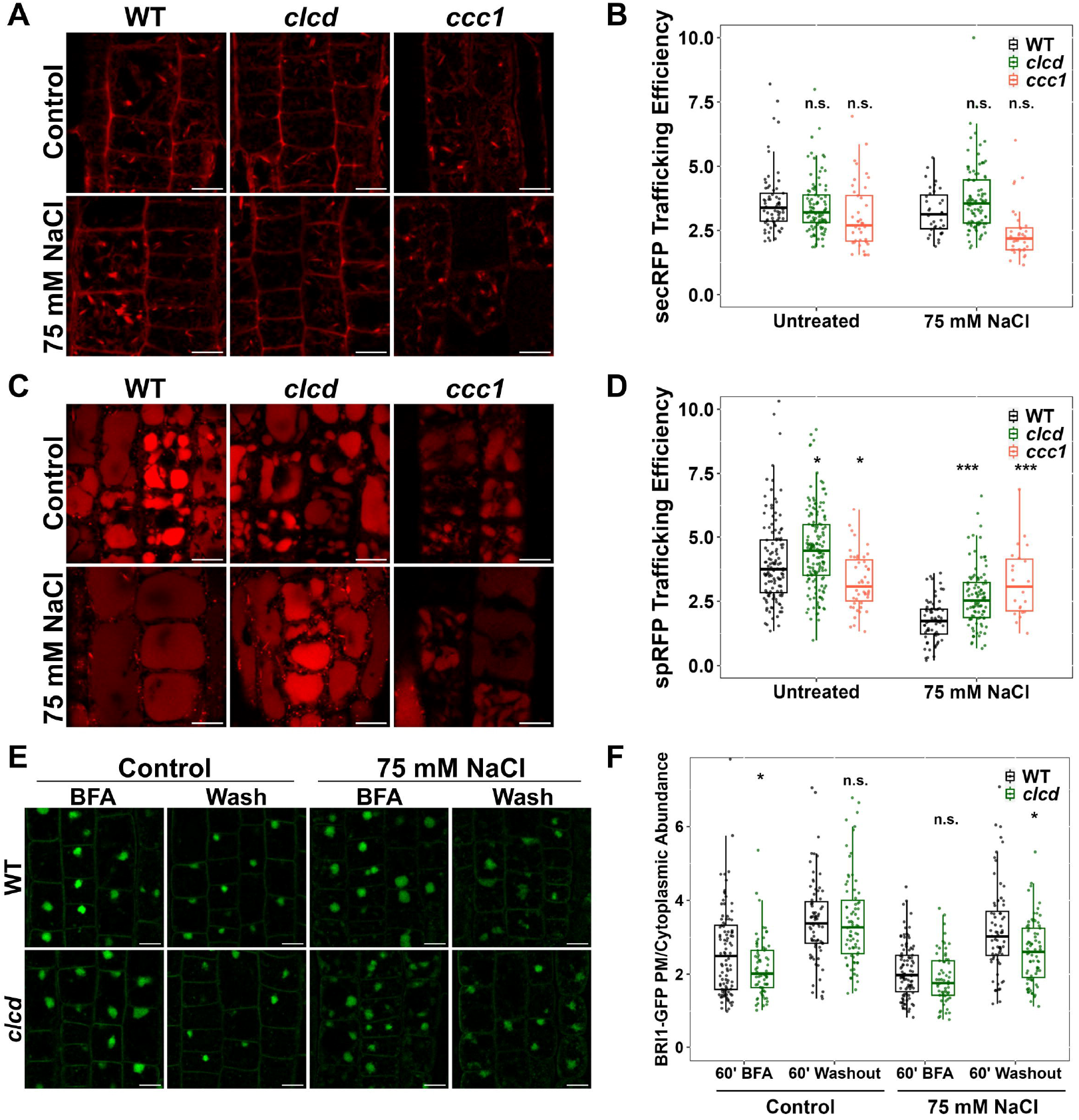
CLCd is not required to maintain bulk cargo trafficking during salt stress. **A-B)** CLSM analysis shows that secretion of secRFP to the apoplast is not affected in *clcd* plants grown on media containing 75 mM. Secretion was measured in root epidermal cells of the elongation zone in 9 day old plants. secRFP trafficking efficiency was determined by measuring the ratio of secRFP signal at the cell periphery relative to the signal in the cell interior. Images are a single slice. Scale bars = 10 um. **C-D)** CLSM analysis shows that secretion of spRFP to the vacuole is not affected in *clcd* plants grown on media containing 75 mM. Trafficking was measured in root epidermal cells of the elongation zone in 9 day old plants. spRFP trafficking efficiency was determined by measuring the ratio of spRFP signal in the vacuole relative to the signal exterior to the vacuole. Images are a single slice Scale bars = 10 um. **E-F)** CLSM analysis of BRI1-GFP recovery after a 1 hour BFA treatment followed by washout and 1 hour recovery shows that clcd plants grown on 75 mM NaCl have a slight reduction in BRI1-GFP recovery compared to WT. A ratio of BRI1-GFP signal at the PM relative to the cell interior is use to determine the relative abundance at the PM where a greater ratio indicates a greater degree of recovery. Imaging was performed in root epidermal cells of the elongation zone of 9 day old plants. Images are single slices. Scale bar = 10 um. For all plots, statistical significance was determined via one-way ANOVA, comparing mutant genotypes to WT under the same conditions. P values are indicated by ***<0.001<**<0.01<*<0.05<n.s. For the boxplots, center line = median, box limits = upper and lower quartiles, whiskers = range to a maximum of 1.5× the interquartile range, points = individual data points.

To analyse membrane protein trafficking, the plasma membrane-localised brassinosteroid-insensitive1 (BRI1) receptor, was investigated. BRI1 transits through the TGN/EE during secretion and undergoes constitutive endocytosis to the TGN/EE, from where it is either targeted to the vacuole for degradation or recycled back to the PM (Dettmer et al., 2006; Geldner et al., 2007; Irani et al., 2012). Previous studies demonstrated that BRI1 recycling and secretion are impaired in trafficking-defective mutants including *det3, nhx5 nhx6, big3*, and *big5* (Dragwidge et al., 2019; Luo et al., 2015; Shang et al., 2024; Xue et al., 2019). To assess BRI1 trafficking dynamics, BRI1-GFP (Geldner et al., 2007) plants were treated with the fungal toxin Brefeldin A (BFA) for one hour which causes TGN/EE aggregation and inhibits TGN/EE-to-PM trafficking, resulting in intracellular BRI1-GFP accumulation in BFA bodies. Following BFA washout, cellular recovery enables BRI1-GFP secretion to resume, allowing quantification of protein delivery rates to the PM. WT plants under control conditions exhibit an increased PM-to-cytoplasmic BRI1-GFP signal ratio, demonstrating protein secretion after BFA washout. Consistent with previous findings that membrane trafficking to the plasma membrane remains unaffected under non-stressed conditions in *clcd* mutants, *clcd* mutants display no difference to WT under control conditions, while BRI1-GFP secretion is slightly reduced when treated with 75 mM NaCl (Figure 4 E-F). This result suggests that salt stress may compromise secretory trafficking of membrane proteins in *clcd* mutants. Collectively, these findings indicate that while bulk cargo trafficking remains largely CLCd-independent under salt stress, trafficking of membrane proteins shows subtle sensitivity to CLCd deficiency during salt treatment.

### Salt stress induces alkalisation of the TGN/EE in an ion transporter dependant manner

The mutants *det3, nhx5 nhx6* and *ccc1* exhibit altered TGN/EE pH and perturbed cargo trafficking (Dragwidge et al., 2019; Luo et al., 2015; McKay et al., 2022). To determine if the altered trafficking of BRI1-GFP in *clcd* could be the result of changes in TGN/EE luminal pH, we interrogated TGN/EE pH using the previously described pH sensor, SYP61-pHusion (Luo et al., 2015). The luminal pH of WT plants grown under control conditions, with 75 mM NaCl or with 150 mM sorbitol was measured. Growing plants on media containing 75 mM NaCl induced an alkalisation of the resting TGN/EE pH. This elevated luminal pH was specific to salt stress and was not observed in plants subjected to osmotic stress with 150 mM sorbitol (Figure 5A).

**Figure 5.**
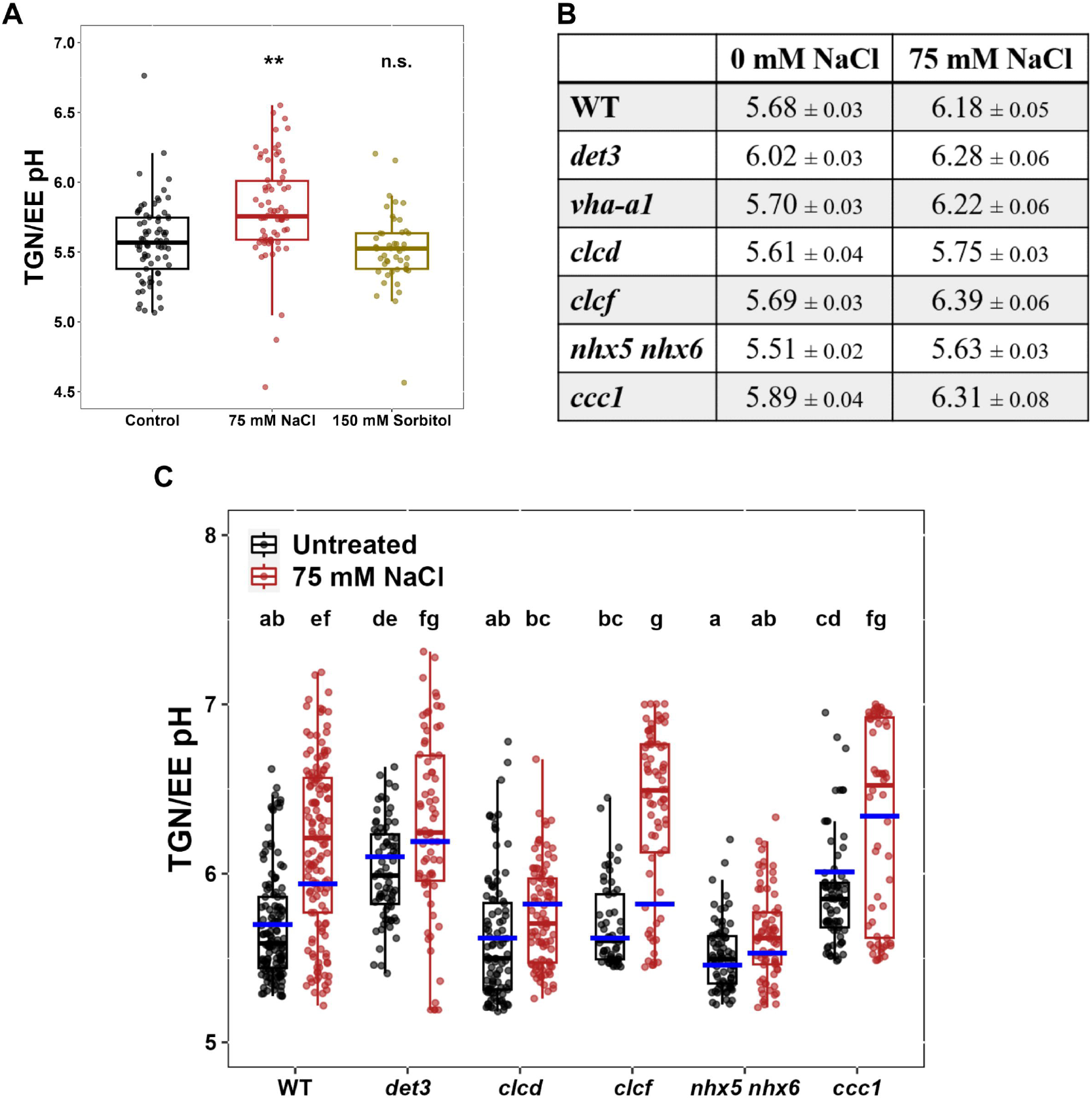
Cytosolic NaCl directly induces TGN/EE alkalisation when CLCd, NHX5 and NHX6 are present. **A)** Measurement of TGN/EE luminal pH in root epidermal cells of the elongation zone using the genetically encoded pH sensor, SYP61-pHusion, reveals that growing WT plants on media containing 75 mM NaCl but not 150 mM Sorbitol results in a more alkaline steady state luminal pH. Statistical significance was determined via one-way ANOVA, comparing to the control condition. P values are indicated by ***<0.001<**<0.01<*<0.05<n.s. **B-C)** Measurement of TGN/EE luminal pH in root epidermal cells of the elongation zone using the genetically encoded pH sensor, SYP61-pHusion, reveals that all genotypes except the Na^+^ hypersensitive genotypes *clcd* and *nhx5/nhx6*, exhibit a TGN/EE pH of at least 6.0 when grown on media containing 75 mM NaCl. Boxplot and points denote the pH measured in plants while the blue bars denote the predicted pH in the computation model. Statistical significance was determined via one-way ANOVA, differing letters denote a P value < 0.05. Printed values = mean ± standard error of the mean.

To determine if the TGN/EE ion transport circuit has a role in salt stress induced TGN/EE alkalisation, the pH of the TGN/EE lumen under control and salt stress conditions was measured in the *det3, clcd, clcf, nhx5 nhx6* and *ccc1* backgrounds. *clcf* plants behaved like WT under control conditions and TGN/EE alkalisation was observed with a 75 mM NaCl treatment (Figure 5B-C). In contrast, the salt hypersensitive *clcd* mutant had a WT-like pH under control conditions but did not exhibit a pronounced alkalisation under salt stress. Compared to WT, the mutants *det3* and *ccc1* both had a higher TGN/EE pH under control conditions and both exhibited only a minor pH increase in the presence of NaCl, resulting in *det3* and *ccc1* having a similar resting pH under salt stress to that of salt stressed WT. The mildly salt hypersensitive *nhx5 nhx6* mutant had a lower TGN/EE pH than WT under control conditions but also did not exhibit a strong increase in pH during salt stress. Together these results demonstrate that the alkalisation of the TGN/EE observed in WT plants under salt stress requires the activity of CLCd, NHX5, and NHX6.

### TGN/EE alkalisation correlates with Na^*+*^ tolerance and VHA-a1 is not limiting for steady-state pH regulation of the TGN/EE during vegetative growth

*clcd, nhx5 nhx6*, and *det3* are Na^+^ hypersensitive but only *clcd* and *nhx5 nhx6* mutants lack the salt induced TGN/EE alkalisation. *det3* mutants further contrast with *clcd* and *nhx5 nhx6* knockouts by exhibiting a sensitivity to the Cl^-^ salts treatment but the *det3* mutation affects V-ATPase activity at both the TGN/EE and the tonoplast. We therefore asked if the Na^+^ hypersensitivity observed in the *det3* mutants is a direct consequence of reduced TGN/EE V-ATPase activity. To address this question, we used a CRISPR-generated knockout allele of the VHA-a1 subunit isoform, *vha-a1-1* (hereafter referred to as *vha-a1*; Lupanga et al., 2020), which mediates TGN/EE localisation of the V-ATPase complex, in order to specifically reduce V-ATPase activity at the TGN/EE without affecting its activity at the tonoplast. Consistent with its WT-like growth during vegetative development, the TGN/EE pH of the *vha-a1* mutant differed only marginally from that of WT under control conditions, suggesting that loss of the VHA-a1 subunit is compensated by VHA-a2 and VHA-a3 isoforms, as previously proposed (Figure 6A; Lupanga et al., 2020). Under salt stress, however, the TGN/EE pH of the *vha-a1* mutant is higher than that of WT under the same conditions, indicating that pH regulation in the TGN/EE of *vha-a1* is compromised under salt stress conditions.

**Figure 6.**
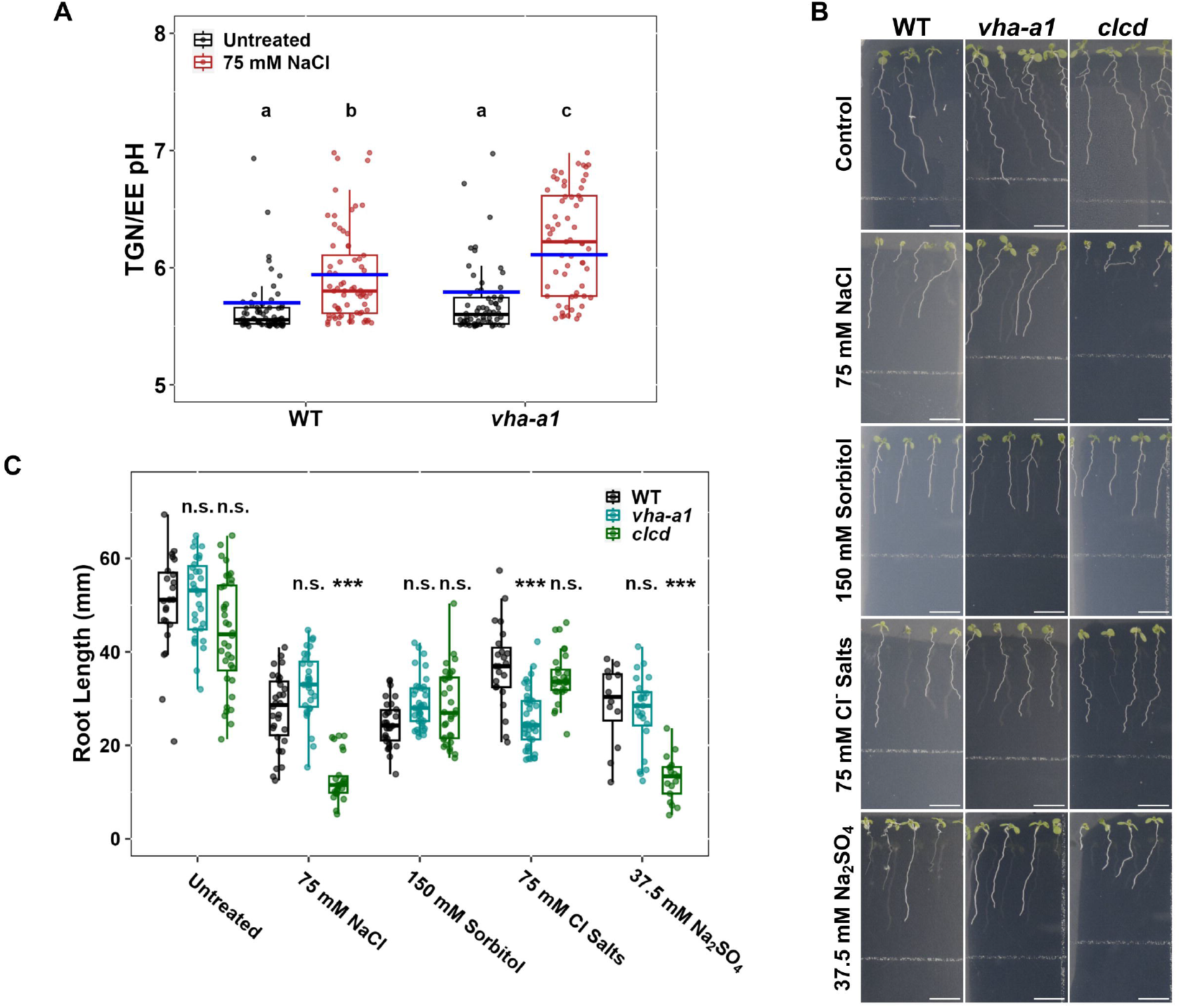
Reducing V-ATPase activity at the TGN/EE alone is insufficient to result in the salt hypersensitivity observed in *det3*. **A)** Measurement of TGN/EE luminal pH in root epidermal cells of the elongation zone using the genetically encoded pH sensor, SYP61-pHusion, shows that *vha-a1* has a higher TGN/EE pH than WT when grown on media containing 75 mM NaCl but not under control conditions. Boxplot and points denote the pH measured in plants while the blue bars denote the predicted pH in the computation model. Statistical significance was determined via one-way ANOVA, differing letters denote a P value < 0.05. Printed values = mean ± standard error of the mean **B-C)** Primary root growth of 14 day old *vha-a1* plants does not exhibit hypersensitivity to Na^+^ exposure. Scale bars = 10 mm. Statistical significance was determined via one-way ANOVA, comparing mutant genotypes to WT under the same conditions. Statistical significance was determined via one-way ANOVA, comparing to the control condition. P values are indicated by ***<0.001<**<0.01<*<0.05<n.s. For the boxplots, center line = median, box limits = upper and lower quartiles, whiskers = range to a maximum of 1.5× the interquartile range, points = individual data points.

Next, we tested whether the compromised pH regulation in *vha-a1* under salt stress is sufficient to cause Na^+^ hypersensitivity by applying a range of salt treatments, including 75 mM NaCl, 150 mM sorbitol, 75 mM Cl^−^ salts, and 37.5 mM Na2SO4. Under NaCl, sorbitol, or Na_2_SO_4_ treatment, no significant difference in root length was observed between *vha-a1* and WT plants, indicating that the TGN/EE-localised VHA-a1 subunit is not required for tolerance to Na^+^ (Figure 6B-C). However, rather than exhibiting Na^+^ sensitivity, *vha-a1* primary root growth showed mild sensitivity to Cl^−^ salts. Taken together, these results demonstrate that VHA-a1 is not required for steady-state pH regulation in the TGN/EE under non-stressed conditions during vegetative growth, but its absence compromises pH regulation under salt stress. Under these conditions, VHA-a1 is not critical for Na^+^ tolerance but appears to contribute to tolerance to Cl^−^ salts. Importantly, the observation that the TGN/EE pH during salt stress is similar in *det3* and *vha-a1* suggests that the elevated TGN/EE pH in *det3* is unlikely to be the direct cause of its Na^+^ hypersensitivity

### TGN/EE pH is directly influenced by cytosolic Na^*+*^ concentrations through the TGN/EE ion transporters

The mutants that lack NaCl-induced TGN/EE alkalisation, *clcd* and *nhx5 nhx6*, are also the Na^+^ hypersensitive mutants suggesting that alkalisation of the TGN/EE may be important during salt stress. We therefore investigated how the alkalisation of the TGN/EE might be achieved during salt stress using mathematical modelling of the TGN/EE ion transport circuit. To build a model reflecting the TGN/EE ion transport circuit in plants, the model’s outputs were compared with TGN/EE pH measurements from WT and knockout mutants. For model construction, no distinction was made between CLCd and CLCf, or between NHX5 and NHX6. To simulate *det3, clcd* and *clcf* plants, V-ATPase or CLC activity was set to 50 % of WT levels, respectively. For *nhx5 nhx6* and *ccc1* knockouts, we initially attempted to model these knockouts with full loss of the respective transport activity. However, it was found that some basal activity was always required to match experimental results, suggesting some redundancy in the system. Thus, we set NHX and CCC activity to 10 % of WT when simulating these knockouts. A model simulating unregulated V-ATPase activity together with CLC, NHX, and CCC transporters, predicted a stable TGN/EE luminal pH of 6.60 for WT, which is significantly higher than the empirically measured pH. Further assessment of this model revealed further discrepancies, with the model predicting a pH of 3.11 when NHX activity was reduced and a pH of 6.10 when CCC activity was reduced, both severe deviations from the trends observed *in planta* (Figure S2A). To explore whether unidentified K^+^ or Cl^-^ channels might contribute to TGN/EE pH regulation, we extended the model to include open K^+^ and Cl^−^ channels. A range of models was tested in which both channel types were present in the same cell or each channel type was present alone. Each configuration was simulated either as low-conductance channels, with ion permeation rates comparable to those of the transporters, or as high-conductance channels, facilitating fluxes far exceeding transporter rates. Nonetheless, none of the tested conditions yielded pH outputs consistent with the empirical values measured across all genotypes in plants (Figure S2A).

Next, we assessed if regulation of protein activity or transporter open probability may be required to achieve the pH values measured in plants. Experimental evidence suggests that CCC1 transport activity may be regulated by luminal pH with Arabidopsis CCC1 activity in oocytes decreasing as extracellular pH decreases from 6 to 5 (Colmenero-Flores et al., 2007). CCC1 activity in the model was therefore placed under the regulation of luminal pH with a linear decrease in activity ranging from 5 % at a luminal pH of 5.4 to 100 % at a luminal pH of 6.0. This change alone was insufficient to simulate measured TGN/EE pH values (Figure S2B).

Further potential changes to protein regulation were therefore investigated, and it was found that the model could successfully recapitulate the pH values measured *in planta* when the following modifications to the model were made: CCC activity was regulated by luminal pH, as described above; V-ATPase activity was regulated by luminal K^+^ concentrations, and NHX activity was modulated by the membrane potential (Figure 5C, blue bars). V-ATPase activity was assumed to decrease linearly with increasing luminal K^+^ concentration from 100 % activity at 250 mM to 5 % activity at 350 mM. Likewise, the model best fit the measured data when NHX exhibited a 2:1 K^+^:H^+^ transport stoichiometry, and its activity linearly depended on membrane potential with 5 % activity at +50 mV (positive inside the lumen) and 100 % activity at −30 mV. It is important to note that, although the modelling results are consistent with pH values observed *in planta*, they do not allow the underlying mechanisms regulating ion transporters to be determined with certainty. Instead, the models indicate the conditions under which we would expect to observe a concurrent change in transporter activity in the biological system. For instance, in the current model, luminal K^+^ concentration is used as the regulatory trait for V-ATPase activity. However, increases in luminal K^+^ are also predicted to correlate with increases in luminal Cl^−^ concentrations, TGN/EE size, and luminal osmolality. Consequently, any one of these traits could produce equivalent modelling outputs and, therefore, might act as a regulator of V-ATPase activity (Figure S2B).

Overall, these additions to the model faithfully reproduce the experimental pH values observed under control conditions for the WT and the various mutants, and it demonstrates that the previously identified TGN/EE ion transporters (i.e. the V-ATPase, CLCs, NHXs and CCC) are sufficient to regulate TGN/EE luminal pH under control conditions in WT plants. To probe the robustness of this system, we assessed the ability of the modelled ion circuit to regulate luminal pH, Cl^-^, K^+^ and TGN/EE size in response to a range of cytosolic pH, Cl^-^ concentrations, K^+^ concentrations, osmolarity, and membrane potential changes. The proposed model was capable of buffering the impact of a variety of different conditions to maintain conditions within the TGN/EE lumen, demonstrating that the proposed system would be sufficiently robust enough to regulate TGN/EE conditions in varying environments and cell types (Figure S3).

To determine how NaCl-induced TGN/EE alkalisation is achieved, it was firstly determined what impact elevated cytosolic NaCl concentrations would have on the TGN/EE ion transport circuit. The addition of 20 mM NaCl to the cytosol was used as a proxy for the effect that 75 mM NaCl applied externally to the root would exert. NHX and CCC were assumed to be Na^+^ permeable with a low selectivity for Na^+^ over K^+^, consistent with experimental results for plant CCC and NHXs (Bassil et al., 2019; Colmenero-Flores et al., 2007). Adding 20 mM NaCl to the cytosol in the model induced an alkalisation of the TGN/EE of a similar magnitude to what was measured *in planta* with the *in planta* luminal pH changing from 5.68 ± 0.03 under control conditions to 6.18 ± 0.05 under salt stress and the model pH shifting from 5.70 under control conditions to 5.94 under salt stress. Furthermore, the model predictions for *det3, clcd, nhx5 nhx6*, and *ccc1* yielded results similar to what was measured in the mutant plants (Figure 5C, blue bars). These results suggest that the TGN/EE alkalisation, which correlates with Na^+^ tolerance, is a direct result of the influence elevated cytosolic NaCl concentrations have on TGN/EE ion transporters. Furthermore, the model revealed that the TGN/EE pH results obtained in the *vha-a1* knockouts, maintaining acidification under control conditions but displaying a higher degree of alkalisation under salt stress, is consistent with a reduction in proton pumping activity of approximately 40 % (Figure 6A, blue bars). This indicates that under control conditions, a 40 % reduction in proton pumping activity at the TGN/EE (*vha-a1*) can be buffered by other TGN/EE ion transporters but not a 50 % reduction (*det3*). To determine if another parameter that could not be measured *in planta*, such as luminal Cl^-^ or K^+^ concentrations, also correlated with Na^+^ hypersensitivity, we determined luminal concentrations of Cl^-^, K^+^ and TGN/EE size under control and salt stress conditions in the model. The assessed parameters did not exhibit NaCl-induced trends specific to the Na^+^ hypersensitive mutants, therefore suggesting that alkalisation, rather than another co-occurring luminal change, is important for Na^+^ tolerance (Figure S4). In summary, through systematic parameter optimization, we successfully developed a mathematical model that faithfully reproduces the coordinated regulation of V-ATPase, CLC, NHX, and CCC transport activity necessary for TGN/EE pH homeostasis under control conditions and accurately recapitulates the empirically observed salt-induced alkalinisation of this compartment.

### TGN/EE pH is influenced by various cytosolic ions in a dose dependent manner

The finding that TGN/EE alkalisation is both a direct response to NaCl and an important aspect for tolerating salt stress, could indicate that this is a potential mechanism for detection of Na^+^ stress. To investigate this possibility, we used the model to determine if the magnitude of the alkalisation has a dose dependant relationship with cytosolic NaCl concentration and if the alkalisation of the TGN/EE is specific to Na^+^ ions. Changes in steady-state pH were assessed while varying cytosolic NaCl concentrations from 0 mM to 50 mM. The TGN/EE pH increased progressively by 0.15 units for every 10 mM NaCl increment. This suggests that the NaCl-induced TGN/EE alkalinization follows a dose-dependent relationship (Figure S5A). To determine if the TGN/EE alkalisation is a result of Na^+^ specifically, the TGN/EE pH was measured in WT plants treated with various Na^+^ and Cl^-^ salts. Specifically, plants were treated with 75 mM NaCl, 75 mM KCl, 75 mM NaNO_3_, or 37.5 mM Na_2_SO_4_. Alkalisation of the TGN/EE was observed in all treatments suggesting that the response is not specific to Na^+^ (Figure S5B). Rather, Cl^-^, K^+^, or Na^+^ all induce alkalisation, although, with differing magnitudes. Supporting these findings *in planta*, our mathematical model also predicts that TGN/EE luminal pH increases in response to different salt additions to the cytosol (Figure S5B, blue bars). Further, modelling the influence of individual ions revealed that Cl^-^ has a greater impact on luminal pH compared to Na^+^ and K^+^, but the final pH change results from of the additive effect of all of the cytosolic ions (Figure S5C).

## Discussion

Here, we find that salt stress induces alkalisation of the TGN/EE lumen in Arabidopsis and that this alkalisation correlates with the plant’s ability to tolerate Na^+^ stress. The finding that cytosolic NaCl can directly influence TGN/EE ion homeostasis and results in alkalisation highlights the importance of endomembrane compartments in crucial cellular adjustments to environmental stresses and the cell’s nutritional state. Sensing of solute concentrations via transporters has previously been theorised (Contador-Álvarez et al., 2025; Dreyer et al., 2022; Wegner et al., 2025). Generally, as solute concentrations change, it subsequently alters the activity of the transporters that bind and transport the solutes, which in turn results in further downstream changes to the membrane potential, pH and Ca^2+^. Such shifts in homeostasis cannot result from the activity of a single transporter but rather, requires a network of transporters, interlinking homeostats and operating as a larger system (Dreyer et al., 2022). Understanding processes such as NaCl mediated TGN/EE alkalisation can therefore only be achieved when treating the ion transport network as a singular system rather than focusing on individual proteins.

In studies focused on single TGN/EE ion transporters, there was a noted association between transport mutants with an altered TGN/EE pH under control conditions and perturbations in vesicular trafficking. *det3, nhx5 nhx6*, and *ccc1* mutants exhibit alterations in TGN/EE pH and have severe developmental phenotypic defects, likely caused by defects in vesicular trafficking (Bassil et al., 2011; Colmenero-Flores et al., 2007; Dettmer et al., 2006; Dragwidge et al., 2019; Luo et al., 2015; McKay et al., 2022; Reguera et al., 2015; Schumacher et al., 1999). Naturally, the most obvious conclusion was that TGN/EE luminal pH was important for vesicular trafficking, especially since *clcd* and *clcf* plants, which have no TGN/EE pH defects, have a WT-like phenotype under control conditions (Scholl et al., 2021). Here, we find that the pH in the TGN/EE lumen of WT plants grown under salt stress is higher than under control conditions but this change in pH does not associate with decreased vesicle trafficking. It is possible that this pH change is required under these conditions to maintain vesicle trafficking although such a conclusion is not supported by the bulk cargo trafficking results in *clcd*, a mutant that does not exhibit TGN/EE alkalisation under salt stress like WT, but maintains bulk cargo trafficking, therefore providing evidence that alkalisation is not required to maintain vesicular trafficking. Rather, it is likely that pH changes in the TGN/EE either induce changes to trafficking of specific cargos, perhaps through alterations to cargo-receptor binding within the TGN/EE, or that the alterations to luminal pH are a symptom of the underlying, but so far unidentified, cause of reduced vesicular trafficking in *det3, nhx5 nhx6*, and *ccc1* plants. Supporting this latter idea, the three mutants all exhibit vastly different phenotypes even while two of these mutants, *det3* and *ccc1*, both possess a more alkaline TGN/EE pH (Bassil et al., 2011; Colmenero-Flores et al., 2007; Dettmer et al., 2006; Dragwidge et al., 2019; Luo et al., 2015; McKay et al., 2022; Reguera et al., 2015; Schumacher et al., 1999). Therefore, the precise role of TGN/EE ion transport on vesicular trafficking within plant cells still needs to be uncovered.

To create a model with outputs aligning with *in planta* TGN/EE pH measurements, certain assumptions had to be made regarding the regulation of transport activity. Specifically, we assumed the activity of the V-ATPase decreased as the TGN/EE size increased, CCC1 activity decreased as TGN/EE luminal pH decreased, and NHX5 and NHX6 transport dropped as the membrane depolarised. These assumptions created a robust, self-regulating model of TGN/EE ion homeostasis and size. Although, it must be noted that for simplification, the TGN/EE was treated as a spherical compartment for modelling. In practice, the size of the TGN/EE is unlikely to change so much as the surface to volume area ratio ergo, as the osmolarity within the TGN/EE increases, the circularity of the TGN/EE is likely to increase resulting in a reduction of available membrane for the creation of vesicles. The system we propose would maintain TGN/EE osmolarity by reacting to decreasing TGN/EE osmolarity by upregulating V-ATPase activity, energising K^+^ and Cl^-^ import through the CLCs and NHXs and increasing the internal osmolarity while the resultant luminal acidification decreases K^+^ and Cl^-^ export through CCC1, shifting the TGN/EE towards a net import of the ions to counteract decreases in luminal osmolarity.

Here, we demonstrate the capacity of intracellular ion transport networks, when functioning as multi-unit systems, to sense environmental stimuli without the need for dedicated ion binding domains or receptor-kinase signalling. Such signalling systems could be present at several cellular membranes enabling perception of various nutrients and toxins in cells. Positioning of ion transport sensors like these at the PM, tonoplast, and in the endomembrane system would further allow cells to comprehensively monitor the status of the cell (endomembrane), the cells external environment (PM) and nutrient/toxin storage (tonoplast). Identifying and characterising transport system-based sensing mechanisms will require further identification of ion transport systems at other membranes, such as the Golgi and ER, and assessment of the abiotic stress tolerance conferred, not by individual transporters, but of whole transport systems.

## Methods

### Plant material and growth conditions

All plants used in this study were *Arabidopsis thaliana* in the Columbia-0 (Col-0) background. This study used the previously described T-DNA insertion lines *clcd* (At5g26240; SALK_042895; Fecht-Bartenbach et al., 2007), *clcf* (At1g55620; SALK_112962; Scholl et al., 2021), *nhx5 nhx6* (At1g54370; GABI_094H05/At1g79610 SALK_100042; Bassil et al., 2011) and *ccc1* (AT1G30450; SALK-145300; Colmenero-Flores et al., 2007). The single nucleotide polymorphism mutant, *det3* (At1g12840; Schumacher et al., 1999) and the *vha-a1* (At2g28520) CRISPR knockout (Lupanga et al., 2020) were also used. The previously described lines CLCd-GFP (Scholl et al., 2021), CLCf-GFP (McKay et al., 2025), NHX6-GFP (Dragwidge et al., 2019), GFP-CCC1 (McKay et al., 2022), secRFP (Zheng et al., 2005), spRFP-AFVY (Hunter et al., 2007), BRI1-GFP (Geldner et al., 2007) and SYP61-pHusion (Luo et al., 2015) were used in this study.

Plants were grown on media containing half strength Murashige and Skoog (1/2 MS; Duchefa Biochemie) media containing 0.5% sucrose, 0.5g/L MES, 0.8% agar adjusted to a pH of 5.6 - 5.8 with KOH. Plants were sown on plates and stratified for at least 2 days at 4°C before being grown vertically at 22°C for 16 hours light, 8 hours dark. The growth duration differed for experiments as specified below.

### Construct design

#### pGGD-GSL(GS linker)-mNeongreen

The mNeongreen sequence was synthesised and amplified with the forward primer, AACAGGTCTCATCAGGTGCTAGCGGTGGCAGCGGTGGCACTAGTGGTGGAGGCGGATCCATGGTGAGCAAGGGCGAGGA and the reverse primer, AACAGGTCTCTGCAGTCACTTGTACAGCTCGTCCA. The PCR product was blunt end cloned into the pJET1.2 vector and verified by sequencing. A verified clone was digested using *Eco31I* to release the PCR fragment. The digested fragment was combined with pGGD00 via green gate cloning.

#### UBQ10:VHA-a1-mNeongreen

pGGC-VHA-a1-intron 10 (Lupanga et al 2020) was used in a GreenGate reaction with the following modules: pGGA006:UBQ10 promoter, pGGB003:N-decoy, pGGD-GSL-mNeongreen, pGGE:HSP18.2 terminator, pGGF007:kanamycin resistance cassette and the destination vector pGGZ004.

#### Salt sensitivity assay

Primary root length was measured to assess the salt sensitivity of plants. Seedlings were grown on media supplemented with an indicated treatment for 14 days. After 14 days, seedlings were imaged with a Nikon digital camera. Images contain white lines present on the equipment used to take the images. Primary root length was measured using FIJI (Schindelin et al., 2012).

#### Protein subcellular Localisation

For determining the localisation of TGN/EE transporters, plants expressing VHAa1-mNeonGreen, CLCd-GFP, CLCf-GFP, NHX6-GFP or GFP-CCC1 were grown for 10 days before imaging. Plants were either grown on standard 1/2 MS media or supplemented with 75 mM NaCl for salt stress. Confocal imaging was performed using a Leica Stellaris 8 microscope with a 63X objective. GFP and mNeongreen were excited at 489 nm and emission was collected between 494-552 nm. All plants were treated with 2µM FM4-64 (Sigma-Aldrich; Excitation = 587, Emission = 610-752) for 10 minutes before imaging to stain the PM. For each construct, 6 plants were imaged. All constructs except for GFP-CCC1 were imaged in the root epidermal cells of the elongation zone. GFP-CCC1 expression is driven by the trichoblast specific promoter, *EXP7*, and therefore can only be imaged in trichoblast cells of the mature root. Following imaging, gaussian blur (radius 0.8) and a background subtraction (radius 50) were performed in FIJI.

#### Protein trafficking

Bulk protein trafficking was measured using the markers secRFP and spRFP (Excitation = 555 nm, Emission = 560-734 nm). Plants expressing the markers were grown for 10 days on standard 1/2 MS media or media supplemented with 75 mM NaCl. The marker was imaged in root epidermal cells of the root elongation zone using a Lecia Stellaris 8 confocal laser scanning microscope with a HC PL APO 63x water immersion objective. Measurement of images was performed in FIJI by first applying a gaussian blur (radius 0.8) and a background subtraction (radius 50) before the average intensity of ER, apoplastic and vacuolar signal was determined for 5 cells per root. A ratio of delivered marker to ER retained marker was then used as a measure of trafficking efficiency.

Trafficking of BRI1-GFP was measured using a BFA washout assay. Plants expressing BRI1-GFP (Excitation = 473 nm, Emission = 478-550 nm) were grown as per the conditions noted for the bulk protein trafficking assay. Seedlings were then treated with brefeldin A (Sigma-Merck) for 1 hour. Half the seedlings were then imaged while the other half were rinsed and transferred to a 1/2 MS liquid solution for 1 hour to washout the BFA before being imaged. Imaging was performed on a Lecia Stellaris 8 confocal laser scanning microscope with a HC PL APO 63x water immersion objective. Trafficking was measured in FIJI by first applying a gaussian blur (radius 0.8) and then a background subtraction (radius 50) before the average intensity of the PM and cell interior signal was determined for 5 cells per root in root epidermal cells of the elongation zone. A ratio of these measurements for each cell was used to determine the efficiency of delivery to the PM.

#### TGN/EE pH Measurements

The pH of the TGN/EE was measured using the SYP61-pHusion sensor as previously described (McKay et al., 2022). Plants expressing this sensor were grown for 10 days on media containing the noted treatments. SYP61-pHusion (RFP Excitation = 555 nm, Emission = 560-734 nm; GFP Excitation = 473 nm, Emission = 478-550 nm) was imaged in root epidermal cells of the elongation zone using a Lecia Stellaris 8 confocal laser scanning microscope with a HC PL APO 63x water immersion objective. A calibration curve was generated using 50 mM MES-BTP solutions containing 50 mM ammonium acetate as previously described (McKay et al., 2022).

#### Computational Modelling

A continuum mathematical model of TGN/EE ion regulation was constructed closely following the work of Ishida and co-workers (Ishida et al., 2013) and previous pH modelling of the secretory pathway of mammalian cells (Wu et al., 2001). The present model tracks the flow of ions and water into/out of the TGN/EE according to the following ordinary differential equations (ODEs):

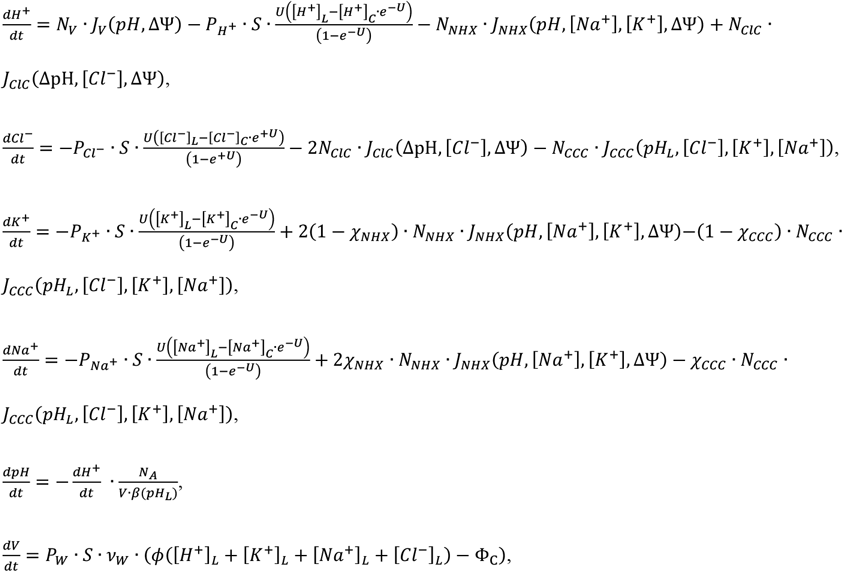

where most symbols are defined in Table 1, *J*_*V*,_ *J*_*ClC*_, *J*_*NHX*_, and *J*_*CCC*_ are the single transport flux rates multiplied by the number of pumps *N*_*V*_, *N*_*ClC*_, *N*_*NHX*_, *and N*_*CCC*_, respectively, *χ*_*NHX*_ and *χ*_*CCC*_ are the Na^+^-to-K^+^ selectivity for the NHX and CCC transporters, respectively, U is the reduced membrane potential, ΔΨ/k_B_T, where k_B_T is a thermal energy unit, and ΔΨ is the membrane potential across the bilayer arising from charge imbalance across the organellar membrane:

**Table 1.** Model parameters.

| description | units | symbol | value | reference |
| --- | --- | --- | --- | --- |
| Cytosolic pH | | $pH_c$ | 7.5 | McKay et al. (2022) |
| Cytosolic potassium concentration | mM | $[K^+]_c$ | 100 | Walker et al. (1996) |
| Cytosolic sodium concentration | mM | $[Na^+]_c$ | 0 / 20 | Ramakrishna et al. (2025), Carden et al. (2003) |
| Cytosolic chloride concentration | mM | $[Cl^+]_c$ | 15 / 35 | Lorenzen et al. (2004) |
| Luminal pH (initial) | | $pH_L$ | 5.7 | Fitted to data |
| Luminal potassium concentration (initial) | mM | $[K^+]_L$ | 284 | Fitted to data |
| Luminal sodium concentration (initial) | mM | $[Na^+]_L$ | 0 | N/A |
| Luminal chloride concentration (initial) | mM | $[Cl^+]_L$ | 165 | Fitted to data |
| Proton permeability | cm/s | $P_{H^+}$ | $6 \times 10^{-5}$ | Grabe (2001) |
| Potassium permeability | cm/s | $P_{K^+}$ | $10^{-7} / 10^{-3} / 0$ | N/A |
| Sodium permeability | cm/s | $P_{Na^+}$ | 0 | N/A |
| Chloride permeability | cm/s | $P_{Cl^-}$ | $10^{-7} / 10^{-3}$ | N/A |
| Bilayer capacitance | $\mu F/cm^2$ | $C_0$ | 1 | Israelachvili (1992) |
| TGN/EE volume | L | $V$ | $4.189 \times 10^{-18}$ | Robinson (2020) |
| TGN/EE surface area | $cm^2$ | $S$ | $1.257 \times 10^{-9}$ | Robinson (2020) |
| Buffering capacity | mM/pH | $\beta$ | 40 | Wu (2001) |
| Osmotic coefficient | | $\phi$ | 0.73 | Grabe (2001) |
| Water permeability | cm/s | $P_W$ | 0.052 | Lencer (1990) |
| Partial molar volume of water | $cm^3/mol$ | $v_W$ | 18 | Dean (1999) |
| Cytoplasmic osmolyte concentration | M | $\Phi_c$ | 1.81 | Argiolas et al. (2016), Miller et al. (2022) |
| Cytoplasmic leaflet charge | mV | $\psi_c^0$ | -30 | Platre et al. (2018) |

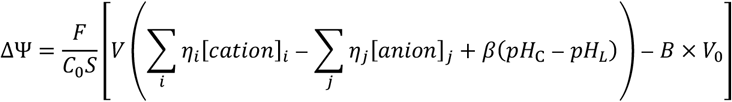

where is F is Faraday’s constant, V is the volume, V_0_ is the initial volume, S is surface area, C_0_ is per area membrane capacitance, i and j run over all cation (except protons) and anion species and their respective valances are η_i_ and η_j_, the term containing the luminal buffering capacity (β) is the total buffered and free protons accumulated as the lumen acidifies from *pH*_*C*_ to *pH*_*L*_, and B is the luminal Donnan particles (sum of impermeant charges). We also consider surface potentials arising from charged lipids on the cytoplasmic side of the membrane 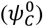 (Platre et al., 2018), and we modify the juxtamembrane cytoplasmic ion concentrations accordingly:

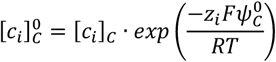

where [*c*_*i*_]_*C*_ is the cytosolic concentration of ion 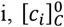 is the value at the TGN/EE membrane surface, and z_i_ is the formal charge. For the flow of ions through channels and transporters, it is the concentration next to the membrane, at the site of entry, that is pertinent, and hence [*c*_*i*_]_*C*,0_ is used in each flux equation. As luminal concentrations change, the surface potentials will also change; however, we assume that the net ionic concentration changes are small enough to ignore this effect. The total membrane potential from bulk cytoplasm to bulk lumen is 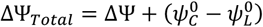, and this value is reported in the manuscript, but the potential difference relevant to the flux of ions across channels and transporters is only ΔΨ.

#### V-ATPase proton pump

The base proton flux 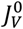 due to the V-ATPase is based on the model in ref. (Grabe et al., 2000) and implemented as in ref. (Grabe et al., 2001). However, we sensitized the turnover rate to luminal K^+^ concentration based on our experiment as follows:

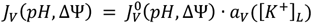

where the activity scales linearly from 1 to 0.05:

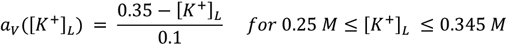

and it remains constant at 1 or 0.05 beyond these concentration bounds.

#### CLC Chloride-Proton antiporter

Our model for the CLC Cl^-^-H^+^ antiporter is taken from the human ClC-7 transporter from ref. (Ishida et al., 2013), which has a 2 Cl^-^ to 1 H^+^ stoichiometry and was fit to experimental data arriving at the following flux:

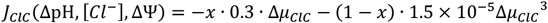

where Δ*μ*_*ClC*_ is the electrochemical potential or driving force, and *x* is a switching function that dictates when the transporter transitions from a linear to a non-linear dependence on the driving force:

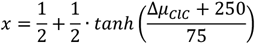

and the electrochemical potential for this transport stoichiometry is:

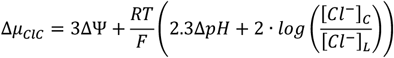

#### NHX Sodium Potassium Proton transporter

We modelled the NHX transport flux as follows:

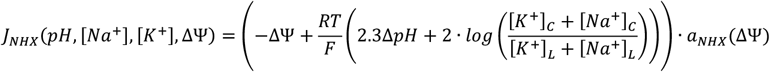

where the activation term a_*NHX*_ linearly decreases from a max value of 1 at ΔΨ = 0 to 0.05 at ΔΨ = ΔΨ_0_= −70 mV and below:

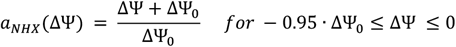

and ΔΨ_0_ = −70 mV, and a_*NHX*_ remains constant for values above and below the range. We assume that the NHX transporter is selective for K^+^ over Na^+^, and the fraction of Na^+^ carried every transport cycle proportional to the fraction of cytoplasmic Na^+^ to the sum of cytoplasmic Na^+^ and K^+^:

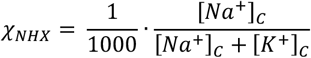

and then the K^+^ flux is then proportional to 1−*χ*_*NHX*_.

#### CCC Chloride Potassium Sodium transporter

The CCC transporter was similarly modelled assuming the transport flux obeys detailed balance and since it is a 1-to-1 anion-cation symporter, there is no dependence on membrane potential:

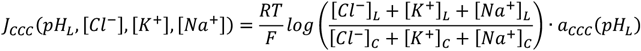

where the activation term a_*CCC*_ linearly decreases from a maximum value of 1 at pH 6 to a minimum value of 0.05 at pH 5.4:

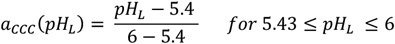

and it remains constant for values above and below the range. Lastly, we assume that the CCC transporter is selective for K^+^ over Na^+^, with the fraction of Na^+^ carried every transport cycle proportional to the luminal fraction of Na^+^ to the total luminal Na^+^ and K^+^ concentration:

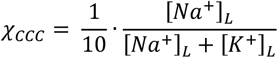

and then the K^+^ flux is then proportional to 1−*χ*_*CCC*_.

## Author Contributions

Conceptualisation, D.W.M, M.K., S.W., K.S.; Formal Analysis, D.W.M, M.U.P.; Funding Acquisition, D.W.M., S.W., K.S.; Investigation, D.W.M., U.L., M.U.P.; Supervision, M.G., K.S.; Writing – original draft, D.W.M, M.G.; Writing – review and editing, D.W.M., U.L., M.K., S.W., M.G.

## Acknowledgements

We thank Jonathon Dragwidge for providing NHX6-GFP, we thank Fabian Fink, Beate Shöfe, Barbara Jesenofsky and Gero Hoffman for assistance with plant and technical work and we thank Ingo Dreyer and Matthew Gilliham for helpful discussion.

## Data Availability

The data underlying this article will be shared on reasonable request to the corresponding author.

## Figure Legends

**Figure S1. CLCd is important for tolerance to Na**^**+**^ **in roots and shoots of mature plants**

The fresh weight of **B)** rosettes and **C)** roots of *clcd* are reduced in comparison to WT and *clcf* when grown for 4 weeks in hydroponics containing 75 mM NaNO_3_ under short day conditions. Scale bars = 10 mm. Statistical significance was determined via one-way ANOVA, comparing mutant genotypes to WT under the same conditions. Statistical significance was determined via one-way ANOVA, comparing to the control condition. P values are indicated by ***<0.001<**<0.01<*<0.05<n.s. For the boxplots, center line = median, box limits = upper and lower quartiles, whiskers = range to a maximum of 1.5× the interquartile range, points = individual data points.

**Figure S2. The V-ATPase, CLCs, NHXs and CCC1 are sufficient to create the TGN/EE luminal pH conditions measured *in planta***. The predicted steady state TGN/EE pH from various modelled scenarios including **A)** without any posttranslational modification of ion transporters (Base Circuit), with the addition of channels for K^+^ or Cl^-^ at transport rates similar to other transporters (Low) or much higher than other transporters (High), **B)** with CCC activity responsive to luminal pH (CCC Regulated) and with V-ATPase activity responsive to various stimuli such as luminal K^+^ concentrations, Cl^-^ concentrations, TGN/EE size (V Responsive) or luminal osmolarity (Osm Responsive). For assessing V-ATPase response to various stimuli, CCC and NHX were both subject to posttranslational modification as described. Magenta lines represent the average measured pH for each genotype as per Figure 4C.

**Figure S3. The modelled TGN/EE ion transport circuit would be sufficient to maintain the ionic homeostasis of the TGN/EE in response to varied external conditions**. The predicted steady state pH when various perturbations to the system are manually introduced shows that the regulated ion transport system proposed for the TGN/EE buffers many cellular changes. The **A)** pH, **B)** luminal Cl^-^, **C)** luminal K^+^ and **D)** TGN/EE size were predicted if the cytosolic pH, Cl^-^ concentration, K^+^ concentration, osmolarity or if the membrane potential were adjusted. ΔMembrane potential represents a manual change to the charge on the cytosolic side of the membrane.

**Figure S4. TGN/EE luminal pH is the only trait found in the mathematical model that correlates with Na**^**+**^ **hypersensitivity in *clcd* and *nhx5 nhx6* plants**. The predicted TGN/EE luminal **A)** Cl^-^ concentration, **B)** K^+^ concentration and **C)** TGN/EE size for each of the ion transport mutants with and without a 20 mM NaCl addition to the cytosol (75 mM NaCl) in our model reveals no pattern aligning with Na^+^ hypersensitivity. TGN/EE size is used as a representation of osmotic adjustment and assumes a perfectly spherical TGN/EE shape.

**Figure S5. TGN/EE alkalisation is a dose dependant response to cytosolic Na**^**+**^, **K**^**+**^ **and Cl**^**-**^. **A)** Measurement of TGN/EE luminal pH in 9 day old WT root epidermal cells of the elongation zone using SYP61-pHusion demonstrates that TGN/EE alkalisation occurs in response to a range of ionic salt treatments. The predicted TGN/EE pH in response to various cytosolic **B)** NaCl concentrations and **C)** ion treatments shows that the ion transport circuit model exhibits TGN/EE alkalisation in a dose dependant manner to Na^+^, Cl^-^ and K^+^. Statistical significance was determined via one-way ANOVA, comparing to the control condition. P values are indicated by ***<0.001<**<0.01<*<0.05<n.s. For the boxplots, center line = median, box limits = upper and lower quartiles, whiskers = range to a maximum of 1.5× the interquartile range, points = individual data points.

