## Supplementary figures and images for "Detection of Cytosolic Ion Concentrations by the Trans-Golgi Network/Early Endosome is Important for Salt Tolerance"

### Supplementary Figure S1

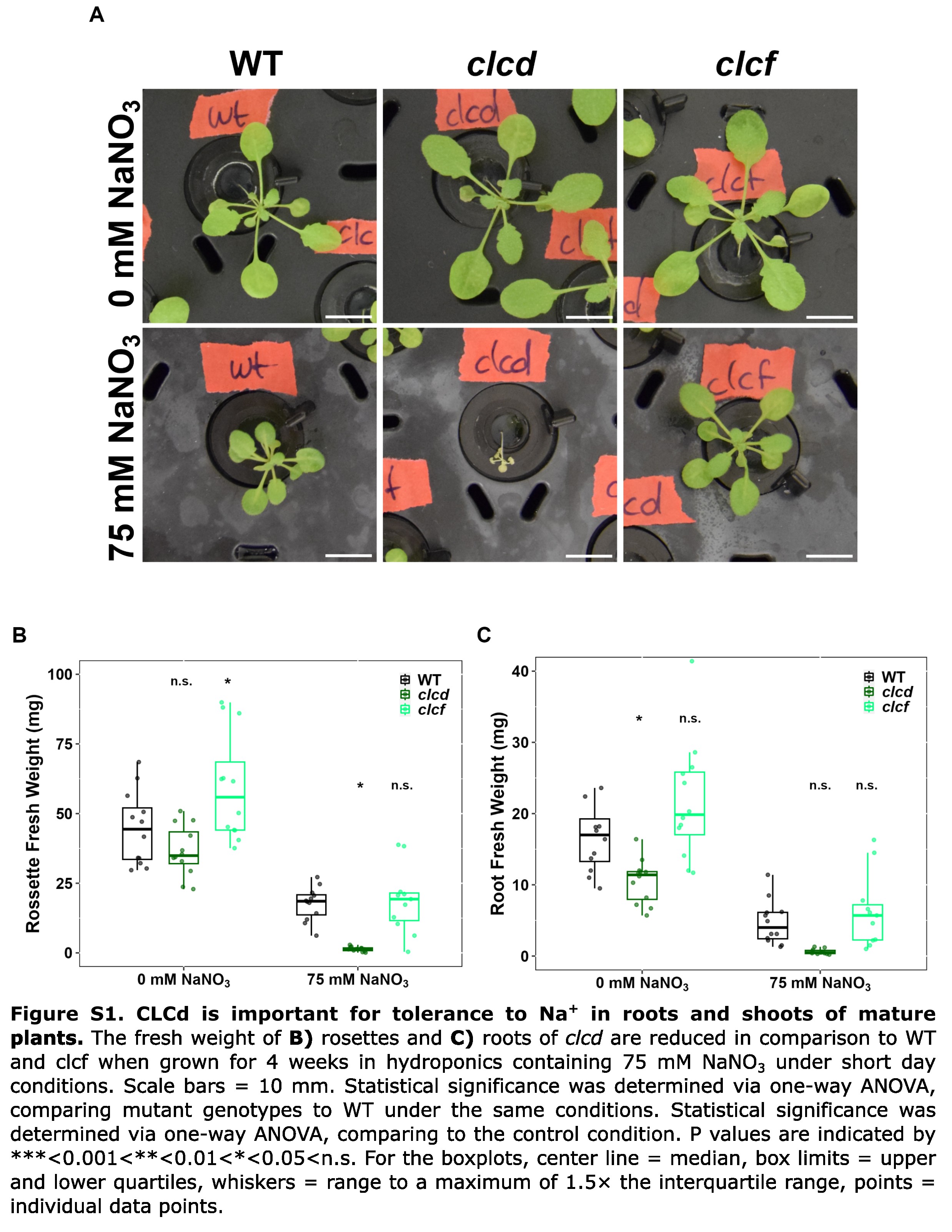

### Supplementary Figure S2

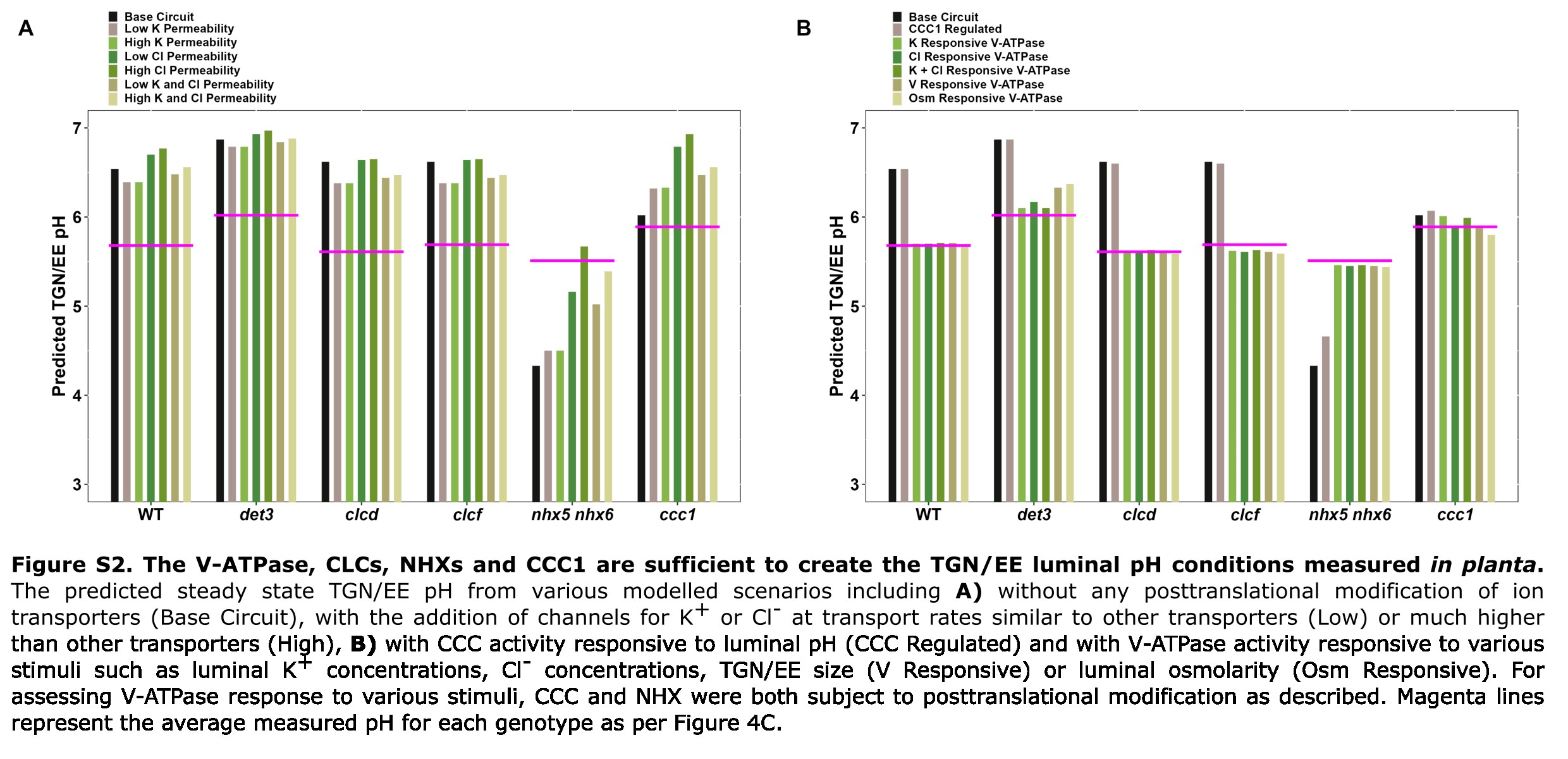

### Supplementary Figure S3

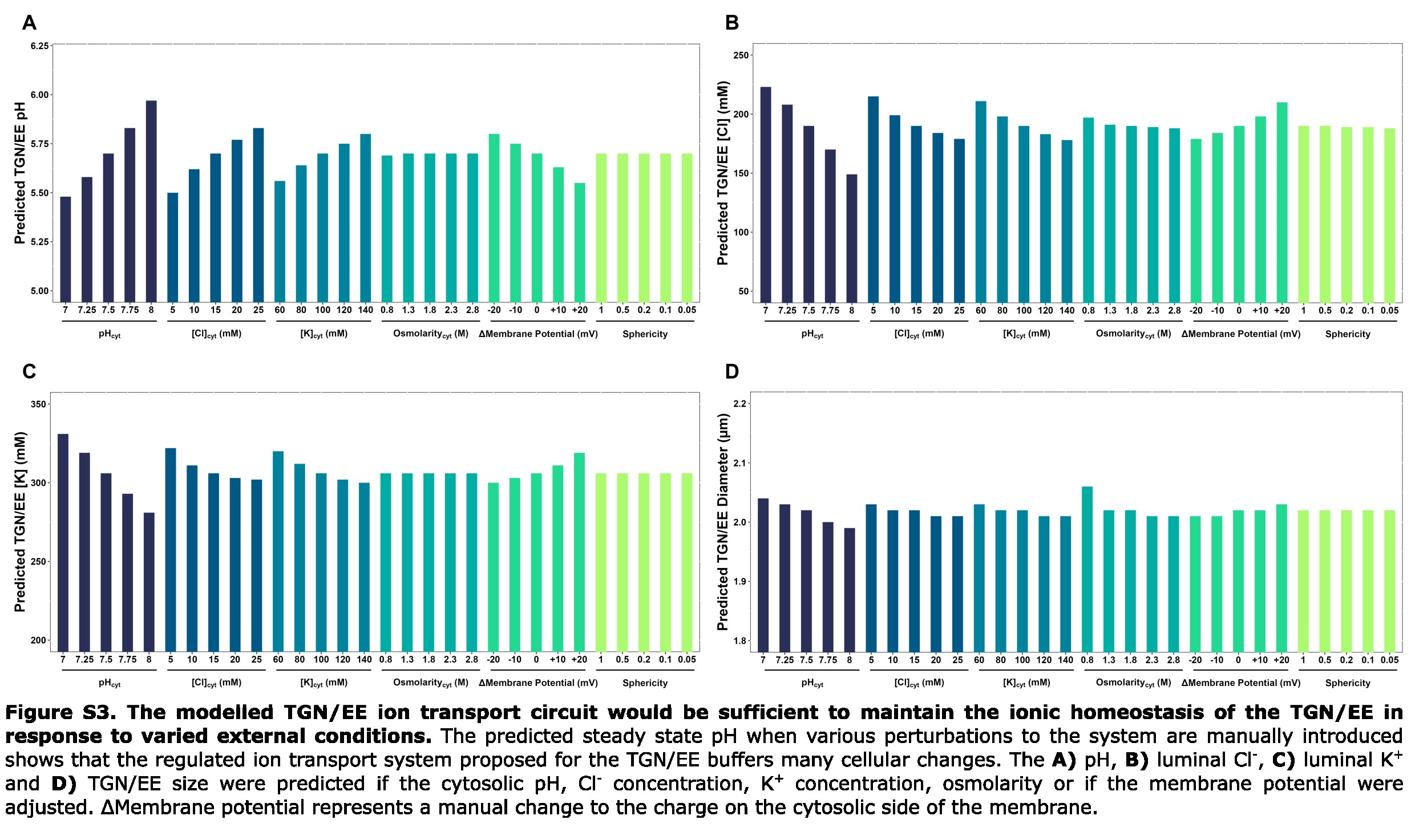

### Supplementary Figure S4

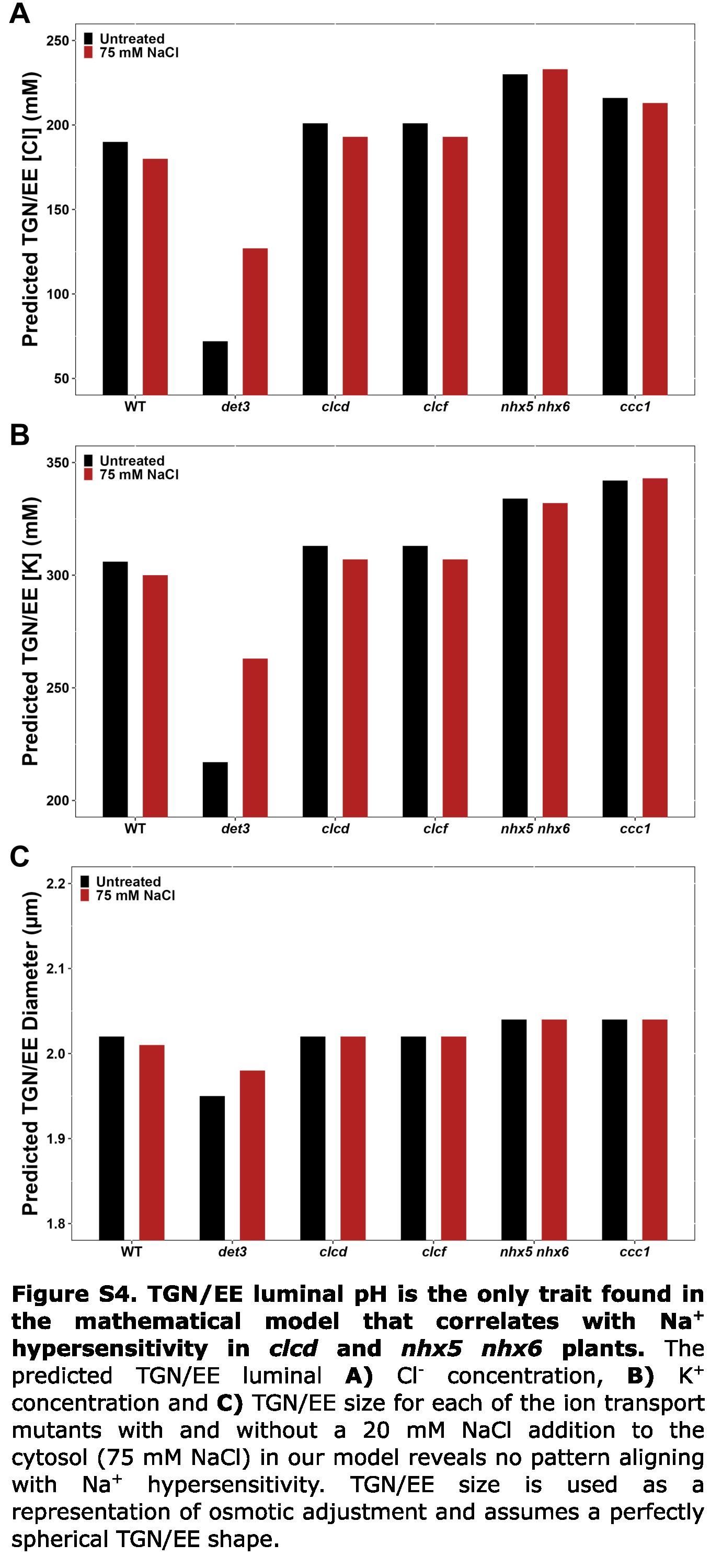

### Supplementary Figure S5

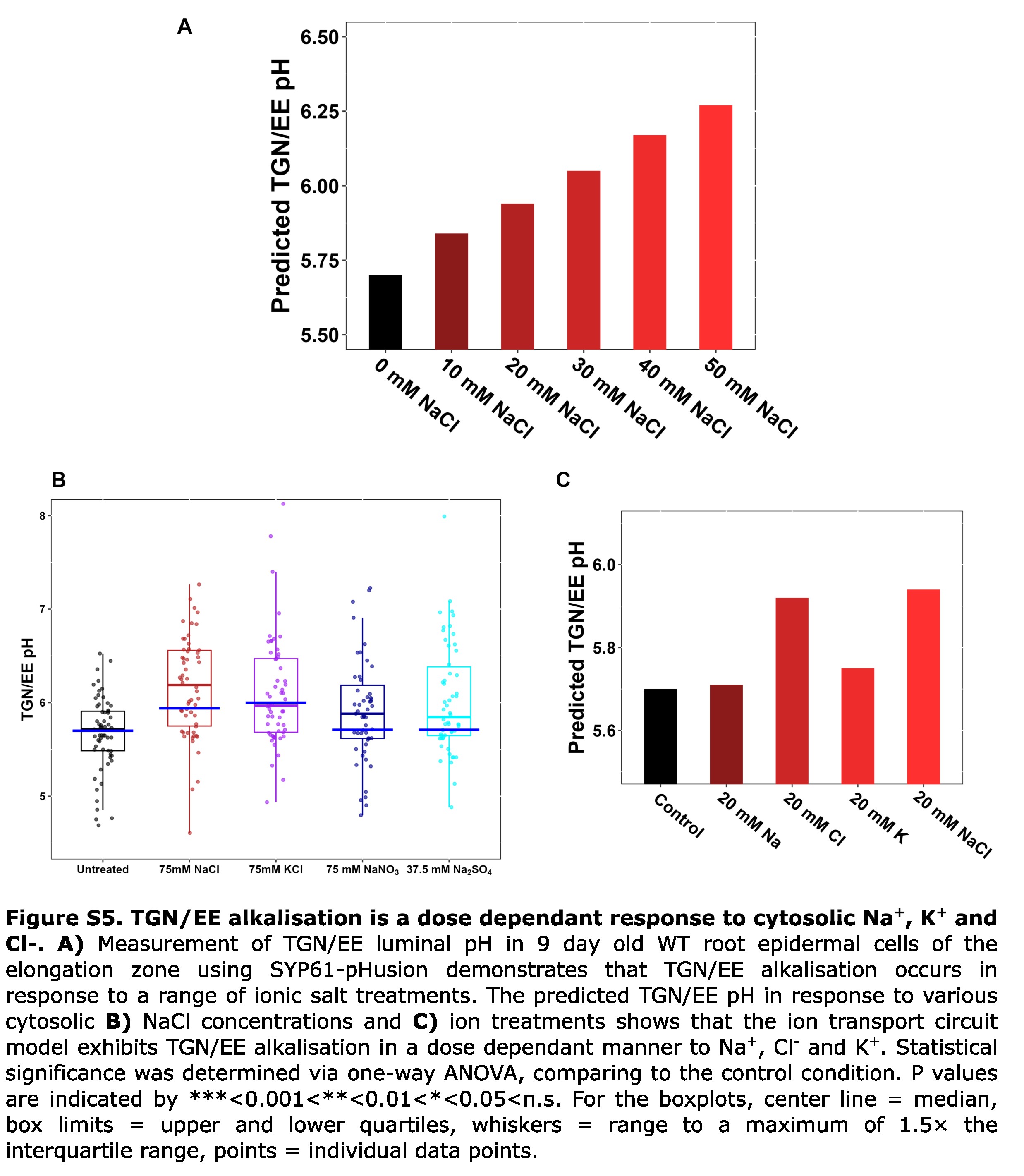
